# Dissecting genomic determinants of positive selection with an evolution-guided regression model

**DOI:** 10.1101/2020.11.24.396762

**Authors:** Yi-Fei Huang

## Abstract

In evolutionary genomics, it is fundamentally important to understand how characteristics of genomic sequences, such as gene expression level, determine the rate of adaptive evolution. While numerous statistical methods, such as the McDonald-Kreitman test, are available to examine the association between genomic features and the rate of adaptation, we currently lack a statistical approach to disentangle the independent effect of a genomic feature from the effects of other correlated genomic features. To address this problem, I present a novel statistical model, the MK regression, which augments the McDonald-Kreitman test with a generalized linear model. Analogous to the classical multiple regression model, the MK regression can analyze multiple genomic features simultaneously to infer the independent effect of a genomic feature, holding constant all other genomic features. Using the MK regression, I identify numerous genomic features driving positive selection in chimpanzees. These features include well-known ones, such as local mutation rate, residue exposure level, tissue specificity, and immune genes, as well as new features not previously reported, such as gene expression level and metabolic genes. In particular, I show that highly expressed genes may have a higher adaptation rate than their weakly expressed counterparts, even though a higher expression level may impose stronger negative selection. Also, I show that metabolic genes may have a higher adaptation rate than their non-metabolic counterparts, possibly due to recent changes in diet in primate evolution. Overall, the MK regression is a powerful approach to elucidate the genomic basis of adaptation.

## Introduction

Understanding the genetic basis of positive selection is a fundamental problem in evolutionary biology. Numerous statistical approaches have been developed to detect loci under positive selection. A popular framework is codon substitution models that seek to infer positively selected genes solely from interspecies sequence divergence (Goldman and Yang, 1994; Muse and Gaut, 1994; Yang *et al*., 2000). By contrasting the rate of nonsynonymous substitutions (*dN*) against the rate of synonymous substitutions (*dS*), codon substitution models can identify positively selected genes with a *dN*/*dS* ratio greater than 1. However, because negative (purifying) selection can dramatically reduce *dN*/*dS* ratios, codon substitution models may be underpowered to detect genes that experienced both positive selection and strong negative selection (Hughes, 2007).

Unlike codon substitution models, the McDonald-Kreitman (MK) test utilizes both interspecies divergence and intraspecies polymorphism to elucidate positive selection in a species of interest (McDonald and Kreitman, 1991; Fay *et al*., 2001; Smith and Eyre-Walker, 2002). By contrasting the levels of divergence and polymorphism at functional sites and putatively neutral sites, the MK test seeks to identify positively selected genes that show an excess of interspecies divergence at functional sites. Because highly deleterious mutations can neither segregate nor reach fixation in a population, the MK test is intrinsically robust to the presence of strong negative selection. On the other hand, weak negative selection may lead to biased results in the MK test because mutations under weak selection can segregate in a population but not reach fixation. To address this problem, several recent studies have extended the MK test to account for the effects of weak negative selection on intraspecies polymorphism (Eyre-Walker and Keightley, 2009; Messer and Petrov, 2013; Galtier, 2016; Haller and Messer, 2017; Uricchio *et al*., 2019), ensuring that the inference of positive selection is not biased by the presence of weak selection. Thus, the MK test and its extensions are powerful methods to disentangle positive selection from ubiquitous negative selection.

Because MK-based methods use relatively sparse divergence and polymorphism data from closely related species, they may be underpowered to pinpoint individual genes under positive selection. To boost statistical power, MK-based methods often are applied to a collection of genes or nucleotide sites with similar genomic features. Using this pooling strategy, previous studies have identified numerous genomic features associated with positive selection in *Drosophila* and primates. The features associated with positive selection in *Drosophila* include local mutation rate (Campos *et al*., 2014; Castellano *et al*., 2016; Rousselle *et al*., 2020), local recombination rate (Marais and Charlesworth, 2003; Campos *et al*., 2014; Castellano *et al*., 2016), gene expression specificity (Fraïsse *et al*., 2019), residue exposure to solvent (Moutinho *et al*., 2019a), X linkage (Avila *et al*., 2015; Campos *et al*., 2018), and sex-biased expression (Pröschel *et al*., 2006; Avila *et al*., 2015; Campos *et al*., 2018). The features associated with positive selection in primates include protein disorder (Afanasyeva *et al*., 2018), virus-host interaction (Enard *et al*., 2016; Uricchio *et al*., 2019), protein-protein interaction (PPI) degree (Luisi *et al*., 2015), and X linkage (Hvilsom *et al*., 2012).

While existing MK-based methods can identify genomic features *associated with* the signatures of positive selection, they may not be able to distinguish genomic features *independently affecting* the rate of adaptive evolution from spurious features *without independent effects* on adaptation (Moutinho *et al*., 2019a; Fraïsse *et al*., 2019). For instance, MK-based methods often are applied to one genomic feature at a time. If a genomic feature with independent effect on the rate of adaptive evolution is strongly correlated with a second feature without independent effect, MK-based methods may report a spurious association between the second feature and adaptive evolution.

Before the current study, two simple heuristic methods have been previously used to estimate the independent effect of a genomic feature on the rate of adaptation by controlling for other potentially correlated features. If we are interested in estimating the independent effect of a gene-level feature, such as tissue specificity, we may estimate the rate of adaption at the gene level and then fit a standard linear regression model, in which we treat the feature of interest and correlated genomic features as covariates and treat the gene-level rate of adaptation as a response variable (Luisi *et al*., 2015; Castellano *et al*., 2016; Moutinho *et al*., 2019a; Fraïsse *et al*., 2019). The regression coefficient associated with the feature of interest can be interpreted as its independent effect on positive selection, holding constant all other genomic features. Although this strategy is powerful and elegant, it cannot be applied to species with low levels of polymorphism, such as primates, due to the challenge of estimating the rate of adaptation at the gene level. Alternatively, we may first stratify genes into a “treatment” group and a “control” group based on the genomic feature of interest. Then, we may use statistical matching algorithms to match each gene from the “treatment” group with a gene of similar characteristics from the “control” group. A significant difference in the rate of adaptation between the two groups of matched genes indicates that the feature of interest has an independent effect on positive selection. Although this method has been successfully used in previous studies (Enard *et al*., 2016; Campos *et al*., 2018; Castellano *et al*., 2019), it is difficult to match genes when there are a large number of genomic features to control for. Therefore, we currently lack a general and powerful statistical framework to estimate the independent effects of genomic features on positive selection by adjusting for a large number of correlated genomic features.

In the current study, I present a novel statistical method, the MK regression, to estimate the independent effects of genomic features on the rate of adaptive evolution. The MK regression is a hybrid of the MK test and the generalized linear regression. Unlike standard linear regression models and statistical matching algorithms, the MK regression can control for a large number of correlated genomic features and is applicable to species with a low level of polymorphism. To the best of my knowledge, the MK regression is the first evolutionary model tailored to characterize the independent effects of genomic features on adaptive evolution. Using synthetic data, I show that the MK regression can unbiasedly estimate the independent effect of a genomic feature even when it is strongly correlated with another genomic feature. Applying the MK regression to polymorphism and divergence data in the chimpanzee lineage, I corroborate previous findings that local mutation rate, residue exposure level, tissue specificity, and immune system genes are key determinants of positive selection in protein-coding genes. In addition, I show that highly expressed genes and metabolic genes may have a higher rate of adaptive evolution than other genes after controlling for several correlated genomic features, which has not been widely reported in previous studies. Taken together, the MK regression is a valuable addition to evolutionary biologists’ arsenal for investigating the genetic basis of adaptation.

## Results

### The MK regression is a generalized linear model tailored to estimate the effects of genomic features on positive selection

The key idea behind the MK regression is to model the site-wise rate of adaptive evolution as a linear combination of local genomic features (Fig. 1). I use *ω*_a_, the relative rate of adaptive substitutions at a functional nucleotide site with respect to the average substitution rate at neutral nucleotide sites, as a measure of the rate of adaptation (Booker *et al*., 2017; Moutinho *et al*., 2019b). Unlike previous MK-based models that treat *ω*_a_ as a gene-level measure, I treat *ω*_a_ as a measure of adaptive evolution at an individual nucleotide site and assume that it can be predicted from local genomic features. To integrate the effects of multiple features on the rate of adaptive evolution, I assume that *ω*_a_, in a site-wise manner, is a linear combination of local genomic features, such as local mutation rate, local recombination rate, and gene expression level. For each genomic feature, the MK regression seeks to estimate a regression coefficient indicating its independent effect on the rate of adaption, holding constant all other genomic features.

**Fig. 1:**
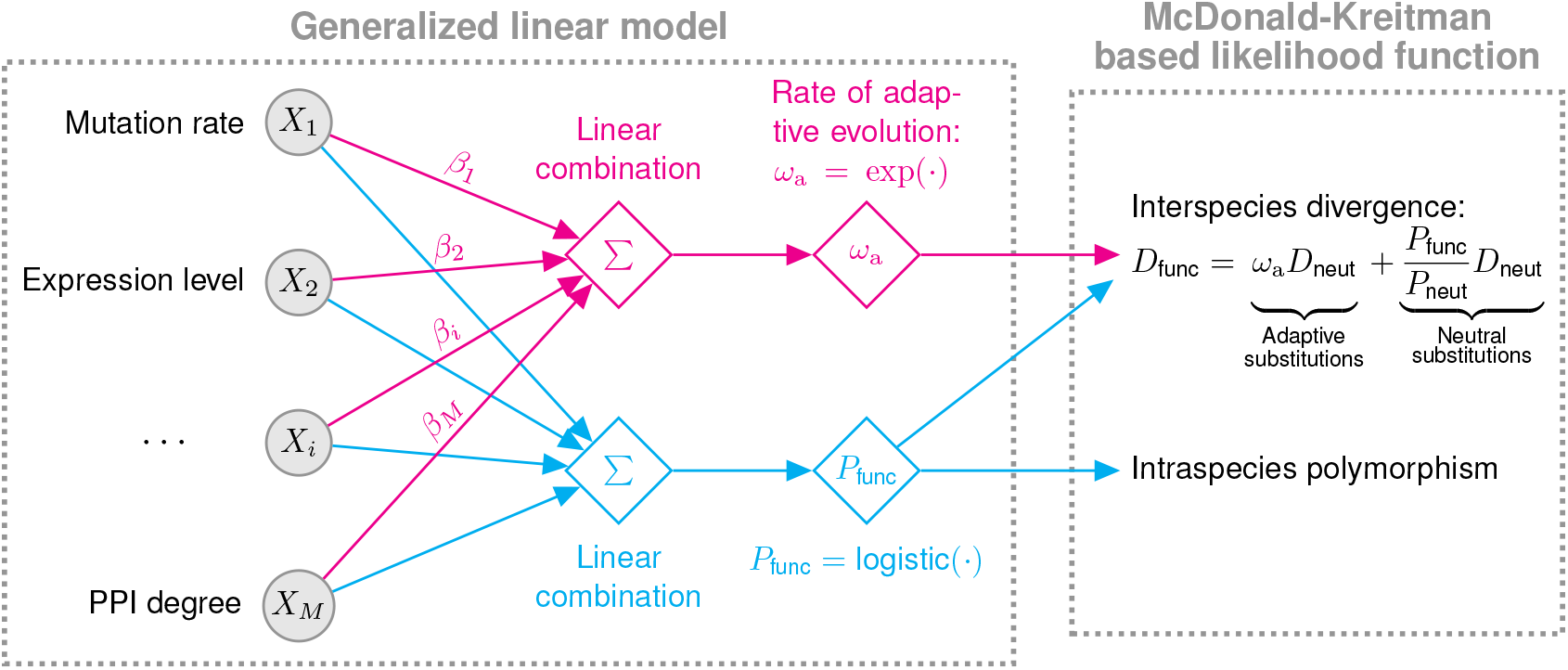
Schematic of the MK regression. The MK regression consists of two components: a generalized linear model and a McDonald-Kreitman based likelihood function. First, I assume that, in a site-wise manner, the rate of adaptive evolution (*ω*_a_) at a functional site is a linear combination of local genomic features followed by an exponential transformation, in which regression coefficient *β*_*i*_ indicates the effect of the *i*-th feature on adaptive evolution. Similarly, I assume that the probability of observing a SNP (*P*_func_) at the same functional site is another linear combination of the same set of genomic features, followed by a logistic transformation. Second, in the McDonald-Kreitman based likelihood function, I combine *ω*_a_ and *P*_func_ at every functional site with two neutral parameters, *D*_neut_ and *P*_neut_, to calculate the probability of observed divergence and polymorphism data given model parameters. *D*_neut_ and *P*_neut_ denote the expected number of substitutions and the probability of observing a SNP at a neutral site, respectively. *D*_func_ denotes the expected number of substitutions at a functional site.

Specifically, the MK regression consists of two components: a generalized linear model and an MK-based likelihood function (Fig. 1). First, I assume that *ω*_a_, in a site-wise manner, is a linear combination of local genomic features followed by an exponential transformation,

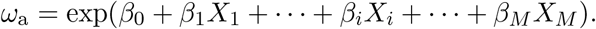

In this equation, *X*_*i*_ is the *i*-th feature at a functional nucleotide site; *β*_0_ is an intercept indicating the baseline rate of adaptive evolution when all genomic features are equal to 0; *β*_*i*_ is a regression coefficient indicating the *i*-th feature’s effect on the rate of adaptation; *M* is the total number of genomic features; exp is an exponential inverse link function which ensures *ω*_a_ is positive. If *β*_*i*_ is statistically different from 0, I consider that feature *i* may have an independent effect on adaptation after adjusting for the other features. Similarly, to accommodate the effects of local genomic features on polymorphism data, I assume that the probability of observing intraspecies polymorphism at a functional nucleotide site, *P*_func_, is another linear combination of local genomic features followed by a logistic transformation (Fig. 1).

Second, in the component of MK-based likelihood function (Fig. 1), I combine *ω*_a_ and *P*_func_ at every functional site with two neutral parameters to calculate the probability of observed polymorphism and divergence data at both functional and neutral sites, which allows for a maximum likelihood estimation of model parameters. Finally, I use the Wald test to examine whether the estimated regression coefficient, 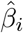, is significantly different from 0 for each feature *i*. It is worth noting that, unlike the standard linear regression, the MK regression does not assume that response variables, *i*.*e*., polymorphism and divergence data, follow a normal distribution. Instead, the likelihood function of the MK regression uses the Jukes-Cantor substitution model (Jukes and Cantor, 1969) and the Bernoulli distribution to describe the generation of divergence and polymorphism in a site-wise manner. Thus, the MK regression can naturally describe the evolution of functional and neutral sites.

### Joint analysis of multiple features distinguishes independent effects from spurious associations

I conducted two simulation experiments to assess the MK regression’s validity and its power to infer the independent effects of genomic features on the rate of adaptive evolution. The simulation experiments consisted of two steps. First, I randomly sampled genomic features from a bivariate normal distribution at each functional site. Second, I generated synthetic polymorphism and divergence data at both functional and neutral sites based on the MK regression model.

In the first simulation experiment, I assumed that there were two genomic features of interest. The first genomic feature had an independent effect on the rate of adaptive evolution, and its regression coefficient, *β*_1_, was equal to 1. On the other hand, the second feature had no independent effect on selection. Thus, its regression coefficient, *β*_2_, was equal to 0 by definition. The other parameters required for the simulation experiment were chosen to ensure that genome-wide levels of polymorphism and divergence are comparable between synthetic data and empirical data from chimpanzees (see details in the Materials and Methods section). To systematically assess the MK regression’s performance with respect to various degrees of correlation between genomic features, I generated four sets of synthetic data with different correlation coefficients between features (0.0, 0.2, 0.4, and 0.6). In each synthetic dataset, I generated 10 independent replicates each of which consisted of 10 Mb functional sites and 10 Mb neutral sites.

I applied two different versions of the MK regression to the synthetic data. The first version was the simple MK regression that analyzed one genomic feature at a time, which was designed to mimic previous MK-based methods. The second one was the multiple MK regression that analyzed two features simultaneously. As shown in Fig. 2A & B, both the simple MK regression and the multiple MK regression produced unbiased estimates of regression coefficients when there was no correlation between features. However, the simple MK regression frequently estimated that 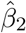 was positive when the two features were correlated with each other, whereas the true value of *β*_2_ was equal to 0. On the other hand, the multiple MK regression always produced unbiased estimates of regression coefficients regardless of the degree of correlation between features.

**Fig. 2:**
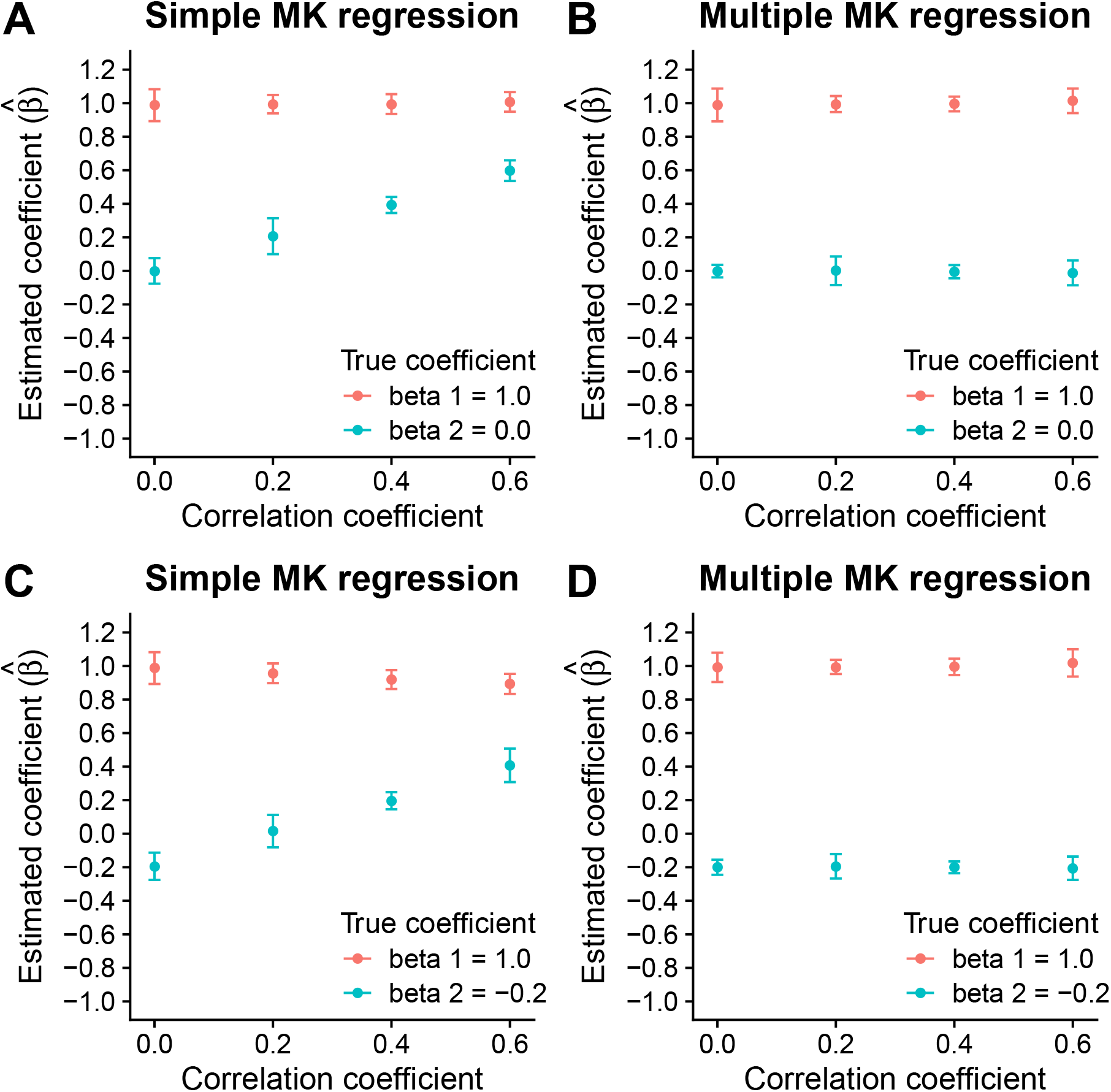
Simulation results. (A) Estimates of regression coefficients in the simple MK regression. The true coefficients are *β*_1_ = 1 and *β*_2_ = 0. (B) Estimates of regression coefficients in the multiple MK regression. The true coefficients are *β*_1_ = 1 and *β*_2_ = 0. (C) Estimates of regression coefficients in the simple MK regression. The true coefficients are *β*_1_ = 1 and *β*_2_ =− 0.2. (D) Estimates of regression coefficients in the multiple MK regression. The true coefficients are *β*_1_ = 1 and *β*_2_ = − 0.2. In each plot, dots and error bars indicate the means and the two-fold standard deviations of estimated coefficients across 10 independent replicates, respectively.

In the second simulation experiment, I evaluated the extent to which the correlation between two causal features complicates the estimation of their independent effects. I set the regression coefficients of the two features to *β*_1_ = 1 and *β*_2_ = −0.2, respectively. Then, I followed the same procedure described in the first simulation experiment to generate synthetic data. As shown in Fig. 2C & D, the multiple MK regression accurately estimated regression coefficients without any noticeable bias, whereas the simple MK regression produced biased results when the two features were correlated with each other. Importantly, when the correlation was strong, the simple MK regression estimated that 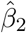 was positive while the true value of *β*_2_ was equal to -0.2.

Furthermore, using the same synthetic data, I evaluated the performance of a previous MK-based method (Smith and Eyre-Walker, 2002; Fraïsse *et al*., 2019), which can only analyze one feature at a time. For each genomic feature, I stratified functional sites into two equal-sized groups. The first group included the top half of functional sites with higher feature value, whereas the second group included the bottom half of functional sites with lower feature value. I estimated *ω*_a_ for each group separately. Then, I calculated Δ*ω*_a_, *i*.*e*., the difference in *ω*_a_ between the two groups of functional sites. If the previous MK-based method can unbiasedly estimate the effects of genomic features, the sign of Δ*ω*_a_ should match the sign of the true regression coefficient. However, the previous MK-based method frequently produced wrong estimates of the sign of Δ*ω*_a_ when genomic features were strongly correlated with each other (Supplementary Fig. 1). In summary, it is critical to jointly analyze multiple genomic features for an unbiased estimation of their independent effects on adaptive evolution.

### The multiple MK regression elucidates genomic determinants of positive selection in chimpanzees

I investigated positive selection in chimpanzee autosomal genes using the MK regression. Because gene annotations were of high quality in the human genome, I converted 4-fold degenerate (4D) and 0-fold degenerate (0D) sites annotated in dbNSFP (Liu *et al*., 2013, 2016) from the human genome to the chimpanzee genome. Because all point mutations at 4D sites are synonymous, I assume that they are putatively neutral. On the other hand, because all point mutations at 0D sites are nonsynonymous, I assume that they are potentially functional. I obtained a genome-wide map of single nucleotide polymorphisms (SNPs) in 18 central chimpanzee (*Pan troglodytes troglodytes*) individuals (de Manuel *et al*., 2016), and inferred ancestral alleles using a reconstructed chimpanzee ancestral genome (Herrero *et al*., 2016; Yates *et al*., 2020). Because the SNP dataset consisted of samples from both females and males, the number of sampled sequences was different between autosomes and sex chromosomes. Thus, I retained only autosomal genes for downstream analysis. To mitigate the impact of weak negative selection on the inference of positive selection, I filtered out SNPs with a derived allele frequency lower than 50% for downstream analysis. In addition, I reconstructed fixed substitutions at 4D and 0D sites in the chimpanzee lineage by comparing the reconstructed ancestral genome with the chimpanzee reference genome. I estimated that the proportion of adaptive nonsynonymous substitutions (*α*) was equal to 15.3% in chimpanzee autosomal genes, which is similar to the estimate in a previous study (Tataru *et al*., 2017).

I collected six genomic features in chimpanzee autosomal genes, including local mutation rate, local recombination rate, residue exposure level, gene expression level, tissue specificity, and the number of unique protein-protein-interaction partners per gene (PPI degree). Specifically, I obtained a chimpanzee-based map of local recombination rates from a previous study (Auton *et al*., 2012) and constructed a map of local mutation rates using putatively neutral substitutions in the chimpanzee lineage. Because functional genomic data were more complete and of higher quality in humans than in chimpanzees, I obtained tissue-based gene expression data from the Human Protein Atlas (Uhlen *et al*., 2015), and utilized the expression level averaged across all tissues and a summary statistic, tau (Yanai *et al*., 2005), as measures of gene expression level and tissue specificity, respectively. I also obtained predicted levels of residue exposure to solvent and experimentally determined PPI degrees in the human genome from previous studies (Wong *et al*., 2011; Luck *et al*., 2020). I converted human-based annotations of gene expression level, tissue specificity, residue exposure level, and PPI degree to the chimpanzee genome (panTro4) using liftOver (Haeussler *et al*., 2019).

I first employed the simple MK regression to analyze the effect of one genomic feature at a time, with no attempt to distinguish independent effects from spurious associations. Because the MK regression used a logarithmic link function for *ω*_a_, I explored if a logarithmic transformation of genomic features can improve model fitting. I found that the logarithmic transformation improved the fitting of the simple MK regression for all features but tissue specificity (Supplementary Table 1). Therefore, I applied the logarithmic transformation to all features except tissue specificity throughout this study. In the simple MK regression, the regression coefficients of local mutation rate, residue exposure level, and PPI degree were significantly higher than 0, whereas the regression coefficient of gene expression level was significantly lower than 0 (Fig. 3A & Supplementary Table 2). On the other hand, local recombination rate and tissue specificity were not significantly associated with the rate of adaptive evolution in the simple MK regression (Fig. 3A & Supplementary Table 2).

**Fig. 3:**
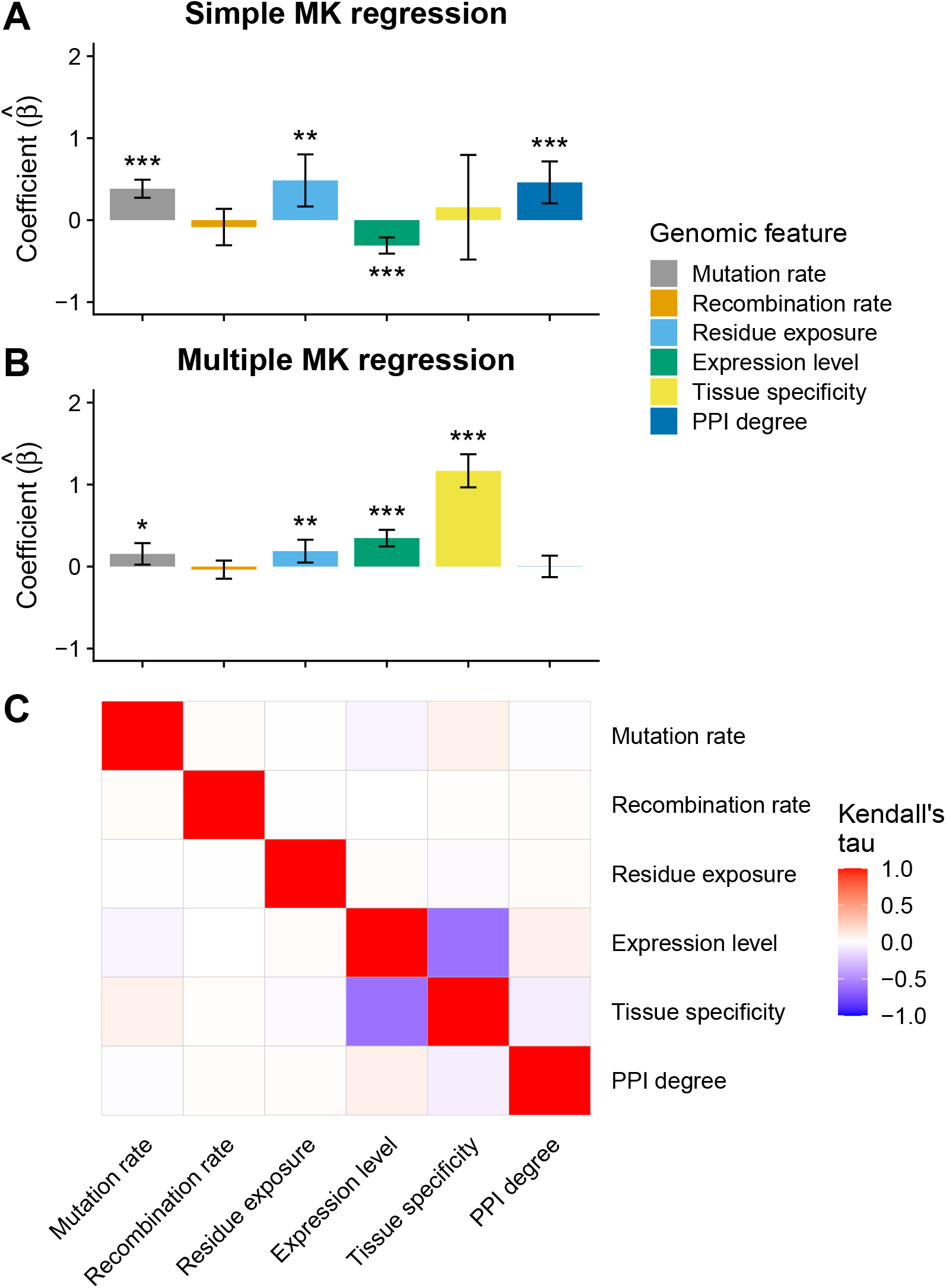
Effects of genomic features on the rate of adaptive evolution. (A) Estimated coefficients of genomic features in the simple MK regression. (B) Estimated coefficients of genomic features in the multiple MK regression. In each bar plot, error bars indicate 95% confidence intervals while one, two, and three asterisks indicate 0.01 ≤ *p*-value < 0.05, ≤ 0.001 *p*-value < 0.01, and *p*-value < 0.001, respectively. (C) Correlations between genomic features.

As discussed in the simulation experiments, the simple MK regression may produce biased estimates of regression coefficients if genomic features are correlated with each other. To test if this was the case in the chimpanzee data, I used the multiple MK regression to analyze the effects of the six genomic features simultaneously. Surprisingly, while local mutation rate, local recombination rate, and residue exposure level showed similar effects in the multiple MK regression, the regression coefficients of the other features were different between the multiple MK regression and the simple MK regression (Fig. 3B & Supplementary Table 3). Specifically, the regression coefficient of PPI degree was not significant in the multiple MK regression (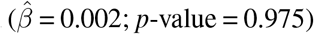, whereas the same coefficient was significant in the simple MK regression 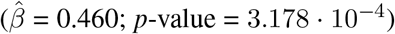. The coefficient of gene expression level was significantly higher than 0 in the multiple MK regression 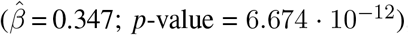, whereas the same coefficient was negative in the simple MK regression 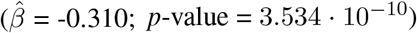. Also, the regression coefficient of tissue specificity was significantly higher than 0 in the multiple MK regression 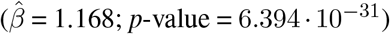 but not in the simple MK regression 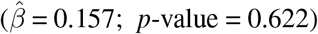.

To examine whether these results were robust to different metrics of tissue specificity, I utilized the negative value of Hg (Kryuchkova-Mostacci and Robinson-Rechavi, 2017) as an alternative metric of tissue specificity. Similar to tau, a higher value of negative Hg indicates a higher level of tissue specificity. I observed qualitatively similar regression coefficients when I replaced tau with negative Hg in the multiple MK regression (Supplementary Fig. 2), although the regression coefficient of local mutation rate was not statistically significant when negative Hg was used. Thus, the estimated effects of genomic features may be robust to different metrics of tissue specificity.

To investigate whether correlations between genomic features could explain the differences in estimated coefficients between the multiple and the simple MK regression, I calculated the Kendall rank correlation coefficient for all pairs of genomic features (Fig. 3C & Supplementary Table 4). I found that local mutation rate, local recombination rate, and residue exposure level were weakly correlated with other features, which may explain why the regression coefficients of these features were consistent between the multiple and the simple MK regression. In contrast, gene expression level and tissue specificity showed a strong negative correlation, which may cause spurious associations in the simple MK regression. I also found that PPI degree was correlated with gene expression level and tissue specificity, although the correlations were to a lesser extent compared with the correlation between gene expression level and tissue specificity. Thus, the observed association of PPI degree with positive selection in the simple MK regression could be due to its correlation with gene expression level and/or tissue specificity.

### Statistical matching analysis confirms genomic determinants identified by the multiple MK regression

I used statistical matching algorithms to corroborate the results of the multiple MK regression. First, I verified whether the association between PPI degree and the rate of adaptive evolution could be explained away by controlling for gene expression level and tissue specificity. I stratified protein-coding genes into two groups with different PPI degrees, in which 1,556 genes with at least 10 protein interaction partners were assigned to the high PPI-degree group whereas 8,471 genes with no more than one interaction partner were assigned to the low PPI-degree group. Without controlling for gene expression level and tissue specificity, the high PPI-degree group had a higher *ω*_a_ than the low PPI-degree group (Supplementary Fig. 3A), but the difference in *ω*_a_ was not significant (*p*-value = 0.146; two-tailed permutation test), possibly due to a reduction of sample size in the stratified analysis. Then, using the default propensity score matching algorithm in MatchIt (Ho *et al*., 2011), I matched each gene from the high PPI-degree group with a gene of similar expression level and tissue specificity from the low PPI-degree group. In the matched data, *ω*_a_ was not different between the high-PPI and low-PPI groups (Supplementary Fig. 3B). I observed similar results using two alternative cutoffs, 5 and 20, for the high PPI-degree group (Supplementary Fig. 4). Thus, PPI degree is unlikely to be a genomic determinant of positive selection in chimpanzees.

I also verified the effect of tissue specificity on the rate of adaptation after adjusting for gene expression level. Due to the strong negative correlation between expression level and tissue specificity (Supplementary Fig. 5), MatchIt returned few matched genes when I attempted to control for expression level. Therefore, I implemented a different matching approach. By closely examining the relationship between expression level and tissue specificity, I found that the variation in tissue specificity was high among highly expressed genes (Supplementary Fig. 5). Thus, I stratified 737 highly expressed genes (gene expression level *>* 30) into two equal-sized groups based on the ranking of their tissue specificity. As shown in Fig. 4A, highly expressed genes with high tissue specificity had a significantly higher *ω*_a_ than their counterparts with low tissue specificity (*p*-value = 0.001; two-tailed permutation test). Therefore, my gene matching analysis confirms the positive effect of tissue specificity on the rate of adaptation in the multiple MK regression.

**Fig. 4:**
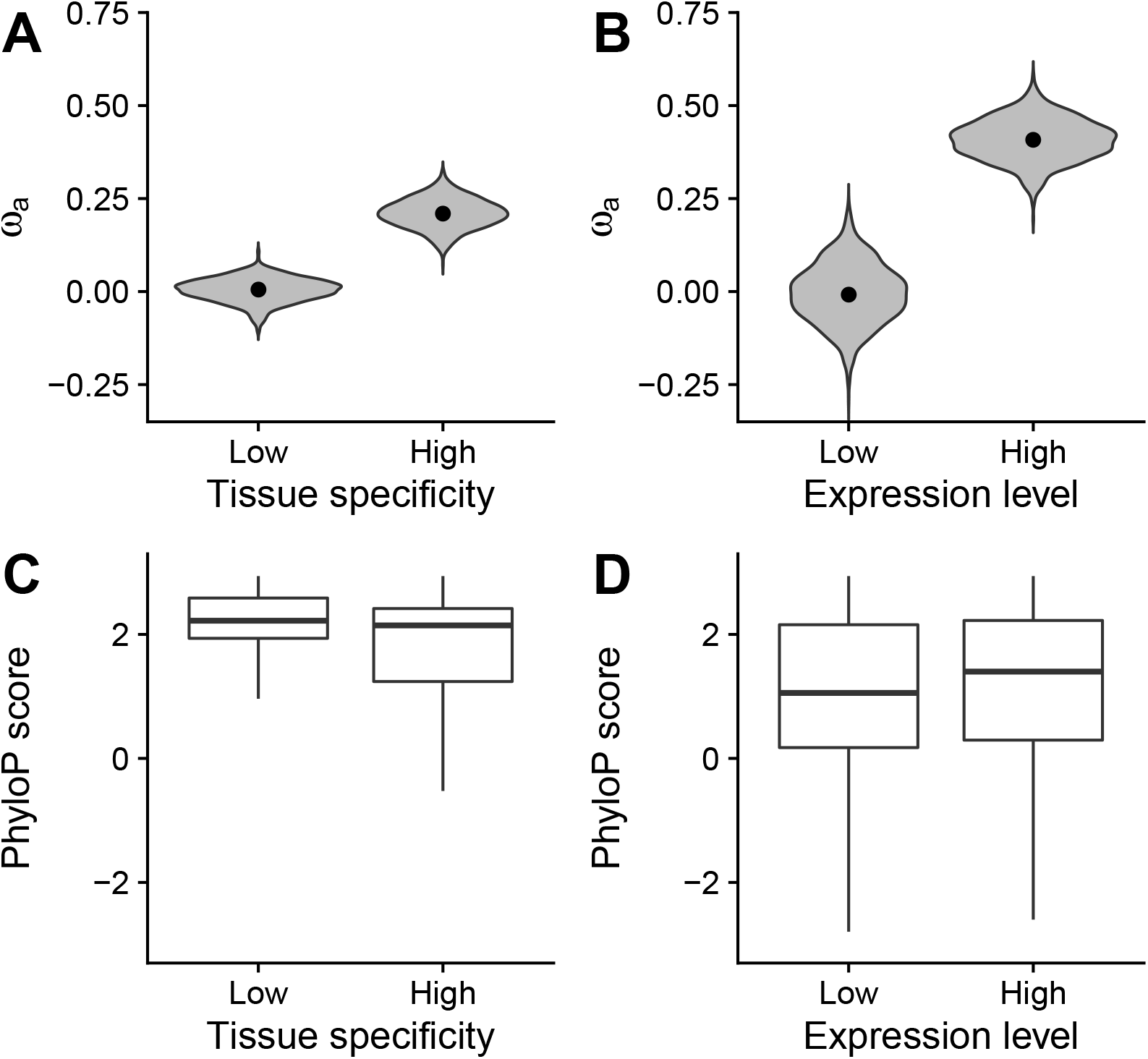
Statistical matching analysis. (A) Estimates of *ω*_a_ in 369 highly expressed genes with low tissue specificity and 368 highly expressed genes with high tissue specificity. (B) Estimates of *ω*_a_ in 498 tissue-specific genes with low expression level and 495 tissue-specific genes with high expression level. In each violin plot, dots indicate point estimates of *ω*_a_ while violins depict the distributions of *ω*_a_ from a gene-based bootstrapping analysis with 1,000 resamplings. (C) Distributions of phyloP scores in 369 highly expressed genes with low tissue specificity and 368 highly expressed genes with high tissue specificity. (D) Distributions of phyloP scores in 498 tissue-specific genes with low expression level and 495 tissue-specific genes with high expression level. In each box plot, the bottom, the top, and the middle horizontal bar of the box indicate the first quartile, the third quartile, and the median of phyloP scores, respectively. The whiskers indicate 1.5-fold interquartile ranges.

As an alternative analysis to infer the independent effect of tissue specificity after controlling for expression level, I divided genes into 10 equal-sized groups (deciles) based on their expression levels. In each decile, I further divided genes into two equal-sized subgroups based on the ranking of their tissue specificity within the decile. As expected, the subgroup with high tissue specificity had a significantly higher *ω*_a_ than the subgroup with low tissue specificity in the decile with the highest expression level (Supplementary Table 5), which confirms the positive effect of tissue specificity on adaptation rate in highly expressed genes. The difference in *ω*_a_ was not significant in other deciles, possibly due to the low variation in tissue specificity in lowly expressed genes (Supplementary Fig. 5).

I observed that the variation in expression level was high among genes with high tissue specificity (Supplementary Fig. 5). To verify the positive effect of gene expression level after adjusting for tissue specificity, I stratified 993 tissue-specific genes (tau > 0.85) into two approximately equal-sized groups based on the ranking of their expression levels. The first group consisted of the top 495 tissue-specific genes with higher expression level, whereas the second group consisted of the bottom 498 tissue-specific genes with lower expression level. The mean expression levels were equal to 5.905 and 0.368 in the two gene groups, which corresponds to a 16-fold difference in mean expression level. As shown in Fig. 4B, tissue-specific genes with high expression level had a significantly higher *ω*_a_ than their lowly expressed counterparts (*p*-value = 0.002; two-tailed permutation test), which confirms the positive effect of gene expression level on the rate of adaptive evolution in the multiple MK regression.

As an alternative analysis to infer the independent effect of gene expression level after controlling for tissue specificity, I divided genes into 10 deciles based on their tissue specificity. In each decile, I further divided genes into two equal-sized subgroups based on the ranking of their expression levels within the decile. As expected, I observed that the subgroup with high expression level had a significantly higher *ω*_a_ than its counterpart in the decile with the highest tissue specificity (Supplementary Table 6). To a lesser extent, *ω*_a_ was slightly lower in the subgroup with high expression level than the subgroup with low expression level in the 7th decile (Supplementary Table 6). The difference in *ω*_a_ was not statistically significant in other deciles, possibly due to the low variation in expression level among genes with low tissue specificity (Supplementary Fig. 5). On average, highly expressed genes showed a higher rate of adaptive evolution than their lowly expressed counterparts.

Also, I used phyloP scores (Pollard *et al*., 2010; Hubisz *et al*., 2011) to examine the effects of gene expression level and tissue specificity on the rate of protein evolution. After controlling for gene expression level, phyloP scores increased with decreasing tissue specificity (Fig. 4C), which is in line with the observation that house-keeping genes tend to evolve at a lower substitution rate than tissue-specific genes (Zhang and Li, 2004; Zhu *et al*., 2008). On the other hand, after controlling for tissue specificity, phyloP scores increased with increasing expression level (Fig. 4D), which is in line with stronger purifying selection on highly expressed genes (Zhang and Yang, 2015). Taken together, it seems that highly expressed genes may be subject to more frequent positive selection than their lowly expressed counterparts, although a higher expression level may impose stronger purifying selection and reduce the overall rate of protein evolution.

### The rate of adaptive evolution may also increase with gene expression level in *Drosophila*

In the previous section, I showed that the rate of adaptive evolution may increase with increasing gene expression level in chimpanzees. Recently, Fraïsse *et al*. (2019) examined the same problem in *Drosophila melanogaster*. Fraïsse *et al*. (2019) first estimated *ω*_a_ for each gene separately. Then, they regressed the gene-level *ω*_a_ on expression level and other potentially correlated genomic features, such as tissue specificity, using the standard linear regression. The coefficients of the standard linear regression were interpreted as the independent effects of genomic features after controlling for other correlated features. Unlike the current study, Fraïsse *et al*. (2019) observed that the rate of adaptation might decrease with increasing expression level in *Drosophila melanogaster*. To reconcile the discrepancy between the current study and Fraïsse *et al*. (2019), I reanalyzed the data from Fraïsse *et al*. (2019). Following Fraïsse *et al*. (2019), I used the standard linear regression to regress the gene-level estimate of *ω*_a_ on gene expression level and tissue specificity, and observed that the coefficient of gene expression was negative (−0.041384; *p*-value = 8.93 · 10^−9^). However, the standard linear regression may suffer from two critical problems in this dataset. First, the residuals of the standard linear regression did not follow a normal distribution, as suggested by a quantile-quantile plot (Supplementary Fig. 6A). Second, the majority of genes had less than 4 nonsynonymous polymorphisms in this dataset (Supplementary Fig. 6B), so the gene-level estimate of *ω*_a_ may be highly inaccurate. Thus, I argue that the standard linear regression may not be an appropriate statistical method for analyzing this dataset.

On the other hand, gene expression level had a positive regression coefficient in the multiple MK regression (Supplementary Fig. 7A). In an orthogonal statistical matching analysis, I stratified 681 *Drosophila* tissue-specific genes (tau > 0.85) into two gene groups based on the ranking of their expression levels. The first group consisted of the top 340 tissue-specific genes with higher expression level, whereas the second group consisted of the bottom 341 tissue-specific genes with lower expression level. Again, tissue-specific genes with higher expression level had a higher *ω*_a_ than tissue-specific genes with lower expression level (Supplementary Fig. 7B) despite that the difference in *ω*_a_ was marginally significant (*p*-value = 0.067; two-tailed permutation test). Taken together, the rate of adaptation may also increase with increasing gene expression level in *Drosophila melanogaster*.

### Metabolic and immune genes may be under frequent positive selection in chimpanzees

To explore whether the positive effect of gene expression level on the rate of adaptive evolution can be explained by the functions of highly expressed genes, I examined the enrichment of Reactome pathways (Jassal *et al*., 2020) and tissue types (Uhlen *et al*., 2015) in the aforementioned 495 chimpanzee tissue-specific genes with high expression level, using the 498 chimpanzee tissue-specific genes with low expression level as a background set. Surprisingly, genes associated with the metabolism of proteins and lipids, and genes with enriched expression in intestine, liver, and pancreas, showed a strong enrichment in the 495 tissue-specific genes with high expression level (Fig. 5A & B; false-discovery rate < 0.01). Based on these results, I hypothesized that metabolic genes may be subject to more frequent positive selection than non-metabolic genes in chimpanzees.

**Fig. 5:**
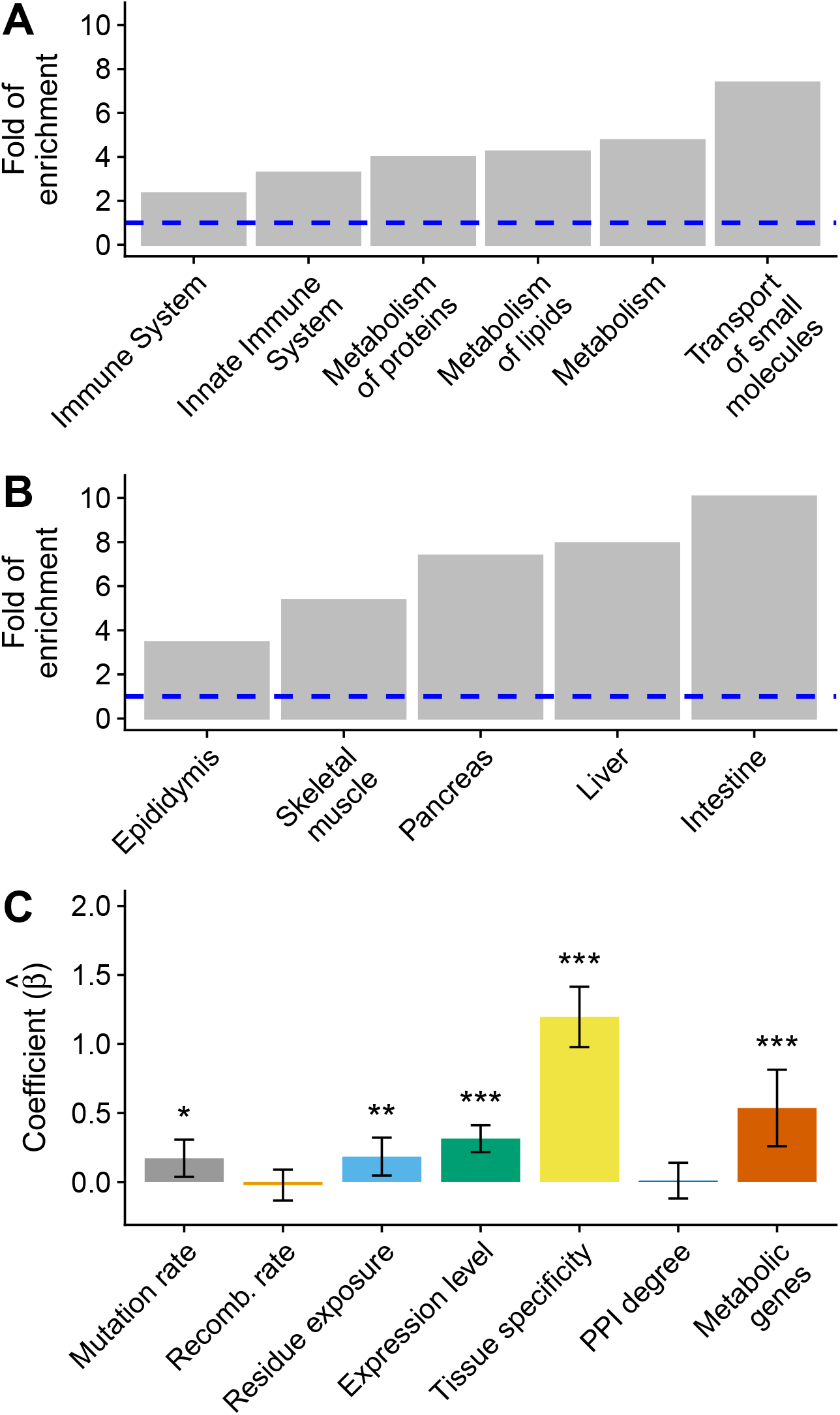
Positive selection in metabolic genes. (A) Enrichment of Reactome pathways in 495 tissue-specific genes with high expression level. (B) Enrichment of tissue types in 495 tissue-specific genes with high expression level. In each enrichment test, 498 tissue-specific genes with low expression level are used as a background gene set. A dashed blue line indicates that the fold of enrichment is equal to 1 (no enrichment). (C) Estimates of regression coefficients in the multiple MK regression. This analysis includes a new binary feature indicating whether each 0D site is located in a metabolic gene. Error bars indicate 95% confidence intervals while one, two, and three asterisks indicate 0.01 ≤ *p*-value < 0.05, 0.001 ≤ *p*-value < 0.01, and *p*-value < 0.001, respectively.

To test this hypothesis, I constructed a genomic feature indicating whether each 0D site was located in one of the 2,220 metabolic genes from MSigDB (Subramanian *et al*., 2005; Liberzon *et al*., 2011). Then, I used the multiple MK regression to simultaneously estimate the effects of the new feature and the six original features. As shown in Fig. 5C & Supplementary Table 7, the regression coefficient of metabolic genes was significantly higher than 0 in the multiple MK regression 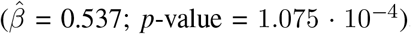, suggesting that metabolic genes might have a higher rate of adaptation than their non-metabolic counterparts. Interestingly, the effect of metabolic genes was not significant in the simple MK regression 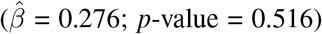. Therefore, controlling for potentially correlated features is critical for revealing elevated positive selection in metabolic genes. Metabolic genes may partially explain the positive effect of gene expression level on adaptive evolution, because the regression coefficient of gene expression level reduced moderately from 0.347 to 0.314 after adding metabolic genes as a new feature in the multiple MK regression (Supplementary Tables 3 & 7).

I observed similar results using propensity score matching. Specifically, I observed that residue exposure level, gene expression level, and tissue specificity showed different distributions between metabolic and non-metabolic genes (Supplementary Fig. 8). Before controlling for these correlated genomic features, *ω*_a_ was similar between metabolic and non-metabolic genes (Supplementary Fig. 8). On the other hand, *ω*_a_ was more than two times higher in metabolic genes than in non-metabolic genes (0.0550 *vs* 0.0234) after controlling for residue exposure level, gene expression level, and tissue specificity (Supplementary Fig. 8) despite that the difference was not statistically significant due to reduced sample size (*p*-value = 0.390; two-tailed permutation test).

To test whether the effect of metabolic genes on adaptation rate can be explained away by their biased expression in digestive organs, I obtained 2,274 genes with biased or enriched expression in digestive organs, including intestine, liver, pancreas, salivary gland, and stomach, from the Human Protein Atlas (Uhlen *et al*., 2015). As expected, metabolic genes were more likely to have biased expression in digestive organs than non-metabolic genes (odds ratio = 2.177; *p*-value < 2.2 · 10^−16^; Fisher’s exact test). Then, I constructed a genomic feature indicating whether each 0D site was located in the 2,274 genes with biased expression. After adding this genomic feature to the multiple MK regression analysis (Supplementary Fig. 9), I observed that the regression coefficient of metabolic genes was still significantly higher than 0 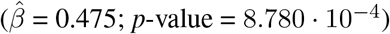, whereas the regression coefficient of digestive system-biased genes was only marginally significant (*p*-value = 0.028). Thus, the effect of metabolic genes on adaptation rate could not be explained by their biased expression in digestive organs.

In line with frequent positive selection on immune system (Schlenke and Begun, 2003; Nielsen *et al*., 2005; Kosiol *et al*., 2008; Barreiro and Quintana-Murci, 2010), I observed that genes associated with immune system had a 2-to 4-fold enrichment in the 495 tissue-specific genes with high expression level (Fig. 5A & B; false-discovery rate < 0.01). To formally test if immune system genes have a higher rate of adaptation than non-immune genes in chimpanzees, I constructed a genomic feature indicating whether each 0D site was located in one of the 3,400 immune system genes from MSigDB (Subramanian *et al*., 2005; Liberzon *et al*., 2011). After adding this new feature to the multiple MK regression analysis, I observed that the regression coefficient of immune genes was significantly higher than 0 (Supplementary Fig. 10 & Supplementary Table 8). Thus, immune genes may have a higher rate of adaptive evolution than their non-immune counterparts. Immune genes may also partially explain the positive effect of gene expression level on the rate of adaptive evolution, since the regression coefficient of gene expression level decreased from 0.314 to 0.238 after adding immune genes as a new feature (Supplementary Tables 7 & 8).

## Discussion

In this work, I have introduced the MK regression, the first evolutionary model for jointly estimating the effects of multiple, potentially correlated genomic features on the rate of adaptive substitutions. Based on similar ideas, my colleagues and I have previously developed statistical approaches to infer negative selection on genetic variants (Huang *et al*., 2017; Huang and Siepel, 2019; Huang, 2020) and the evolutionary turnover of cis-regulatory elements (Dukler *et al*., 2020). Thus, unifying generalized linear models and evolutionary models may be a powerful strategy to address a variety of statistical problems in evolutionary biology.

As shown in the simulation experiments (Fig. 2), when two genomic features are correlated with each other, even at a moderate level, the simple MK regression and a previous MK-based method (Smith and Eyre-Walker, 2002; Fraïsse *et al*., 2019) cannot accurately estimate the independent effects of genomic features because they cannot control for correlated genomic features. On the other hand, the multiple MK regression can unbiasedly estimate the independent effects of genomic features if all relevant genomic features are included in the same analysis. Because we are almost always interested in the independent effects of genomic features, the multiple MK regression may be superior to the simple MK regression and other MK-based methods that can only analyze one feature at a time.

Regression coefficients in the multiple MK regression might be interpreted as the direct causal effects of genomic features on the rate of adaptation. However, similar to other linear regression models (Pearl *et al*., 2016), the causal interpretation of the multiple MK regression relies on two implicit assumptions. First, genomic features of interest and all correlated genomic features should be included in the same MK regression analysis. Second, there should be no reverse causality, *i*.*e*., the rate of adaptive evolution should not cause changes in genomic features. As discussed in the literature of causal inference (Pearl *et al*., 2016), these assumptions cannot be verified using observational data alone and, thus, have to be justified by domain knowledge on a case-by-case basis.

It is worth noting that I do not attempt to infer total causal effects of genomic features on adaptation rate in the current study. According to the theory of causal inference (Pearl *et al*., 2016), the total effect of a genomic feature of interest includes its direct effect on the rate of adaptation as well as its indirect effect through mediators, *i*.*e*., others genomic features that reside on a directed path from the genomic feature of interest to the rate of adaptation in an assumed causal graph. Thus, inferring the total effect of the genomic feature of interest will require me to impose very strong assumptions about the causal relationship between genomic features (see examples in Laubach *et al*. 2021 and Rosenbaum *et al*. 2020). On the other hand, to infer the direct effect of the genomic feature of interest, I can simply control for all potentially correlated features without specifying their causal relationship with the genomic feature of interest (Laubach *et al*., 2021). In other words, the MK regression effectively assumes a simplified causal graph where I do not specify the directions of causality between genomic features (Supplementary Fig. 11; see also Chapter 4.3 in Shipley 2016).

It is also worth noting that I have ignored potential interactions between genomic features in the current study. It is possible to add interaction terms to the multiple MK regression. However, because of the large number of potential interaction terms and the spareness of polymorphisms in the chimpanzee genome, I may lack statistical power to detect interactions between features in chimpanzees.

Using the multiple MK regression, I have identified numerous genomic features with independent effects on adaptive evolution in the chimpanzee lineage (Fig. 3B). First, in line with previous studies (Campos *et al*., 2014; Castellano *et al*., 2016; Rousselle *et al*., 2020), I have shown that the rate of adaptation increases with increasing mutation rate. Because mutations are the ultimate source of genetic variation, a higher mutation rate will increase the genetic variation for positive selection to act on.

Second, previous studies have shown that local recombination rate is positively correlated with the rate of adaptation in *Drosophila* (Marais and Charlesworth, 2003; Campos *et al*., 2014; Castellano *et al*., 2016), probably due to a reduced effect of Hill-Robertson interference in recombination hotspots. However, I have not observed the same pattern in chimpanzees, which could be explained by a reduced impact of linked selection in species with a small census population size, such as chimpanzees (Corbett-Detig *et al*., 2015). Alternatively, my analysis may have limited power to detect a weak association between recombination rate and positive selection due to the lower level of polymorphism and the smaller proportion of adaptive substitutions in chimpanzees compared with *Drosophila* (Castellano *et al*., 2016).

Third, in agreement with a previous study (Moutinho *et al*., 2019a), I have shown that the rate of adaptive evolution increases with the increasing level of residue exposure to solvent. On the other hand, it is well-known that the site-wise rate of protein evolution is positively correlated with the level of residue exposure, possibly due to relaxed negative selection on exposed residues (Goldman *et al*., 1998; Franzosa and Xia, 2009; Liberles *et al*., 2012; Echave *et al*., 2016). Taken together, exposed residues on protein surfaces may be subject to both weaker negative selection and more frequent positive selection than their buried counterparts. From a biophysical perspective, missense mutations on protein surfaces are less likely to disrupt protein stabilities than mutations in hydrophobic cores (Bloom *et al*., 2005, 2006a,b; Franzosa and Xia, 2009). Thus, missense mutations on protein surfaces may be under more frequent positive selection because they are less likely to perturb protein folding. Alternatively, protein surfaces may have a higher rate of adaptive evolution because they may play an important role in host-pathogen interactions (Moutinho *et al*., 2019a).

Fourth, I have shown that the tissue specificity of a gene has a positive effect on the rate of adaptation after controlling for correlated genomic features, such gene expression level. Thus, tissue-specific genes are more likely to be under positive selection than housekeeping genes. Because nonsynonymous mutations in housekeeping genes have a higher chance to disrupt multiple phenotypes, my findings support that the pleiotropic effect is a key determinant of adaptive evolution (Fraïsse *et al*., 2019).

Fifth, I have shown that the rate of adaptive evolution increases with increasing gene expression level in both chimpanzees (Fig. 3B & 4B) and *Drosophila* (Supplementary Fig. 7) after controlling for correlated genomic features, such as tissue specificity. Recently, Fraïsse *et al*. (2019) reported an opposite trend in *Drosophila* by regressing a gene-level estimate of *ω*_a_ on gene expression level and potentially correlated features. However, my reanalysis of their data suggests that the standard linear regression used in Fraïsse *et al*. (2019) may not be an appropriate method for inferring the effects of genomic features in *Drosophila*. Unlike the standard linear regression, the MK regression does not rely on inaccurate estimates of *ω*_a_ at the gene level, and does not assume that the response variables follow a normal distribution. Thus, the MK regression may be more broadly applicable than the standard linear regression in inferring the effects of genomic features on adaptation.

Sixth, numerous studies have shown that immune genes may have a higher rate of adaptive evolution than other genes in various species (Schlenke and Begun, 2003; Nielsen *et al*., 2005; Kosiol *et al*., 2008; Barreiro and Quintana-Murci, 2010). Using the MK regression, I have observed the same trend in chimpanzees (Supplementary Fig. 10 & Supplementary Table 8). Thus, immune genes may be subject to constant adaptation in chimpanzees to fight against ever-evolving pathogens and parasites.

Last but not least, I have shown that highly expressed genes are more likely to be associated with metabolic pathways and digestive organs than their lowly expressed counterparts (Fig. 5A & B), which implies that frequent positive selection in metabolic genes may partially explain the positive effect of gene expression level on the rate of adaptation. In agreement with this hypothesis, I have shown that metabolic genes may have a higher rate of adaptation than their non-metabolic counterparts after controlling for potentially correlated genomic features (Fig. 5C). Similarly, a recent study has reported that metabolic pathways may be subject to more frequent positive selection than non-metabolic pathways in multiple inner branches of the primate phylogeny (Daub *et al*., 2017). Taken together, genes in metabolic pathways may be subject to frequent positive selection in multiple primate species, possibly due to recent changes in diet in primate evolution (Daub *et al*., 2017; Haygood *et al*., 2007; Blekhman *et al*., 2008, 2014). In future studies, it is also interesting to examine whether other genomic features related to metabolism, such as the replication timing of genes (Chen *et al*., 2010) in digestive organs, have independent effects on the rate of adaptive evolution using the MK regression.

Frequent positive selection in metabolic genes has not been widely reported in primates, except in the current study and in Daub *et al*. (2017). I speculate that the discrepancy could be explained by the unique design of the MK regression and the method in Daub *et al*. (2017). First, previous studies focused on identifying individual genes with significant signals of positive selection. If metabolic pathways are under polygenic selection, the signal of selection in a single gene may be too weak to reach genome-wide significance (Csilléry *et al*., 2018; Barghi *et al*., 2020). In contrast, the MK regression and the method in Daub *et al*. (2017) have pooled data across a large number of metabolic genes, which may significantly increase the statistical power to detect diffused signals of polygenic selection (Barghi *et al*., 2020). Second, if metabolic genes are under lineage-specific adaptation in primates, positive selection may only be detected by statistical approaches tailored for a single branch of the primate phylogeny, such as the MK regression and the branch-site codon substitution model (Daub *et al*., 2017). Third, many previous methods may not be able to control for the effects of correlated genomic features. In the current study, I have shown that the effect of metabolic genes is manifested in the multiple MK regression but not in the simple MK regression, which highlights the importance of controlling for correlated genomic features. In future studies, it is tempting to test these hypotheses for a comprehensive understanding of adaptive evolution in metabolic genes.

Similar to a previous study (Luisi *et al*., 2015), I have found that the rate of adaptation increases with increasing PPI degree in the simple MK regression (Fig. 3A). However, the same pattern cannot be replicated in the multiple MK regression (Fig. 3B). Also, after controlling for gene expression level and tissue specificity using propensity score matching, I have found no difference in the rate of adaptation between genes with high PPI degree and genes with low PPI degree (Supplementary Fig. 3B). Thus, PPI degree is unlikely to be a key determinant of positive selection in chimpanzees.

Because functional genomic data are scarce in the chimpanzee genome, I have mapped multiple genomic features from the human genome to the chimpanzee genome to examine adaptive evolution in the chimpanzee lineage. It is worth noting that the current study does not require or assume that these genomic features are perfectly correlated between humans and chimpanzees. Unlike studies that aim to identify individual loci under positive selection, I focus on examining genome-wide relationships between genomic features and adaptation rate. As long as genomic features are well correlated between humans and chimpanzees at the genome-wide scale, my results should be robust to species differences in genomic features in a small set of genes. Because of the short divergence time between humans and chimpanzees, I expect that human features are reasonable proxies of corresponding chimpanzee features at the genome-wide scale. Nevertheless, it is of interest to revisit the results reported in the current study when chimpanzee-specific annotations become available in the future.

While the MK regression is a powerful framework to estimate the effects of multiple genomic features simultaneously, it has a few limitations that are worth of future exploration. Notably, the MK regression is based on the classical MK test and, thus, inherits its limitations (McDonald and Kreitman, 1991; Smith and Eyre-Walker, 2002). First, the MK regression does not explicitly model the effects of weak negative selection on polymorphism data. To mitigate this problem, I have used a simple strategy to filter out low frequency SNPs. This strategy may not be optimal, because a large proportion of SNPs cannot be used in the MK regression despite the fact that they are potentially informative of positive selection (Messer and Petrov, 2013). Second, the MK regression assumes that positively selected mutations have a negligible contribution to polymorphisms. This assumption is valid when weak positive selection is rare. However, there is evidence for frequent weak positive selection in the human genome, and ignoring the effects of weak positive selection may lead to an underestimation of adaptation rate in humans (Uricchio *et al*., 2019). Thus, the MK regression may not be suitable for examining positive selection in the human genome. Third, the MK regression does not explicitly model the impact of demography on polymorphisms. In future studies, it is tempting to extend the MK regression by explicitly modeling the effects of weak negative selection, weak positive selection, and demography on the site-frequency spectrum of polymorphisms (Eyre-Walker and Keightley, 2009; Messer and Petrov, 2013; Galtier, 2016; Haller and Messer, 2017; Uricchio *et al*., 2019).

From a statistical point of view, several aspects of the MK regression may also be improved in future studies. First, it is tempting to relax the strong assumption of a linear relationship between genomic features and the rate of adaptation. For instance, we may replace the generalized linear model in the MK regression by a generalized additive model (Hastie, 1990), which can accommodate more complicated relationships between features and selection while maintaining the interpretability of the MK regression. Second, the MK regression is mainly designed to infer the effects of genomic features on the rate of adaptive evolution. Thus, it may not be the best tool for pinpointing individual genes under frequent positive selection. If positive selection at the gene level is of the main interest, and if the numbers of polymorphic and divergent sites are large in a gene, the classical MK test (McDonald and Kreitman, 1991; Smith and Eyre-Walker, 2002) may be more appropriate for inferring positive selection at the gene level. Third, the MK regression currently can only accommodate the fixed effects of genomic features. It is tempting to extend the MK regression by introducing a gene-level random effect (Huang, 2020), which may allow for estimating the rate of adaptation at the gene level. Fourth, the MK regression currently can only estimate the effects of genomic features on *ω*_a_. Based on the extensive literature on the MK test (Booker *et al*., 2017; Moutinho *et al*., 2019b), it is possible to extend the MK regression to estimate the effects of genomic features on other measures of natural selection, such as the rate of nonadaptive evolution (*ω*_na_) and the proportion of adaptive substitutions (*α*). Fifth, similar to other multiple regression models, the MK regression might yield unreliable estimates of regression coefficients if there is strong collinearity between genomic features (Dormann *et al*., 2013). The current study is unlikely to be affected by strong collinearity because the absolute values of correlation coefficients between genomic features are smaller than 0.7 (Supplementary Table 4), an established cutoff for strong collinearity (Dormann *et al*., 2013). Nevertheless, in future studies, it is tempting to explore rigorous statistical techniques, such as the variance inflation factor (Fox and Monette, 1992), to detect and handle potentially strong collinearity in the MK regression. Finally, when used as a part of a causal inference pipeline, the MK regression is not an automatic tool and requires domain knowledges of all genomic features being considered in an analysis. I expect that future extensions of the MK regression will enable systematic explorations of the genomic basis of adaptive and nonadaptive evolution in various species, such as humans and *Drosophila*.

## Materials and Methods

### Details of the MK regression

The MK regression consists of two components: a generalized linear model and an MK-based likelihood function. In the generalized linear model, I assume that 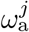, the relative rate of adaptive substitutions at functional site *j*, is a linear combination of genomic features followed by an exponential transformation,

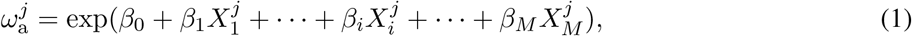

in which 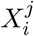 is the *i*-th local genomic feature at site *j, β*_*i*_ is a regression coefficient indicating the effect of feature *i* on 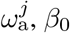 in an intercept, and *M* is the total number of genomic features. Because genomic features may also influence the levels of polymorphism at functional sites, I model the probability of observing a SNP at functional site *j*, 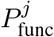, as another linear combination of genomic features followed by a logistic transformation,

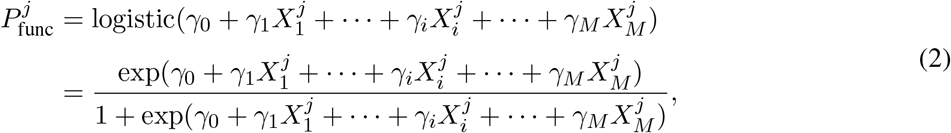

in which *γ*_0_ and *γ*_*i*_ are an intercept and a regression coefficient with respect to 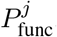, respectively. Similar to the regression coefficients in equation 1, *γ*_*i*_ represents the effect of feature *i* on the occurrence of polymorphism at functional site *j*. It is worth noting that I effectively assume an infinite-site model here (Kimura, 1969), so no more than one SNP is allowed at a single site. Finally, I introduce two neutral parameters, *D*_neut_ and *P*_neut_, which represent the expected number of substitutions and the probability of observing a SNP at a neutral site, respectively. These neutral parameters are shared by all neutral sites.

In the MK-based likelihood function, I specify the probability of polymorphism and divergence data at both neutral and functional sites given model parameters (*β*_0_, *β*_*i*_, *γ*_0_, *γ*_*i*_, *D*_neut_, and *P*_neut_). First, I denote 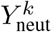 as a binary response variable indicating the presence/absence of a SNP at neutral site *k* and assume that it follows a Bernoulli distribution,

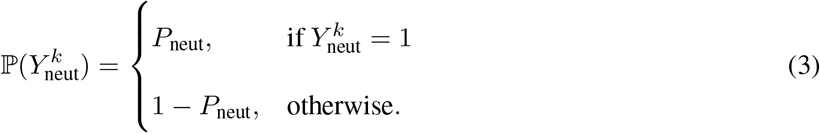

Similarly, I denote 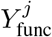 as a binary response variable indicating the presence/absence of a SNP at functional site *j* and assume that it follows a Bernoulli distribution,

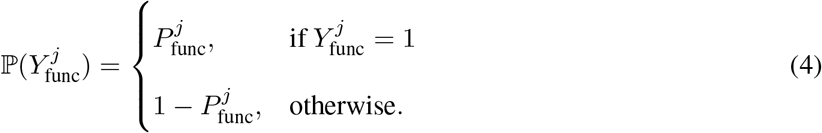

Second, I employ the Jukes-Cantor substitution model (Jukes and Cantor, 1969) to describe the distribution of interspecies divergence at neutral sites. Denoting 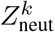 as a binary response variable indicating if the reference genome and the ancestral genome have different nucleotides at neutral site *k*, the Jukes-Cantor model suggests that

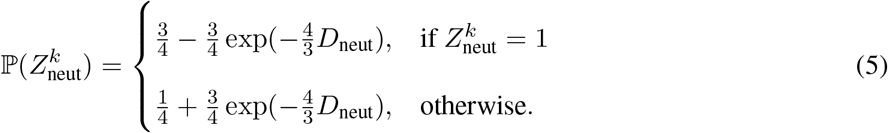

Third, to model the effects of positive selection and neutral factors on interspecies divergence at functional sites, I assume that 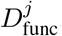, *i*.*e*., the expected number of substitutions at functional site *j*, is equal to the sum of the number of adaptive substitutions and the number of neutral substitutions (Bierne and Eyre-Walker, 2004; Gossmann *et al*., 2010),

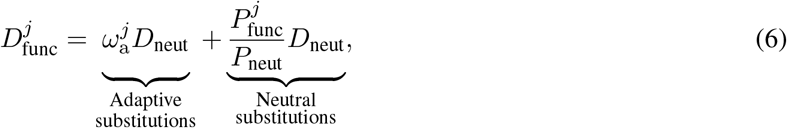

in which 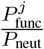 is equal to the relative rate of neutral evolution at functional site *j* compared with neutral sites. Given 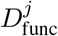 at each functional site, I again employ the Jukes-Cantor substitution model to describe interspecies divergence at functional site *j*,

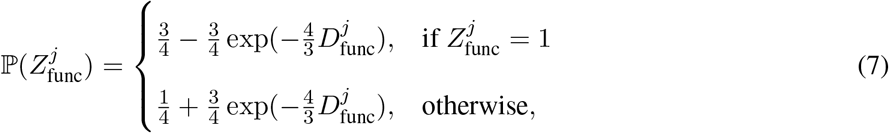

in which 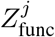 indicates if the reference genome and the ancestral genome have different nucleotides at functional site *j*. Finally, I assume that nucleotide sites evolve independently given genomic features and model parameters. Thus, I define the MK-based likelihood function of the whole dataset as

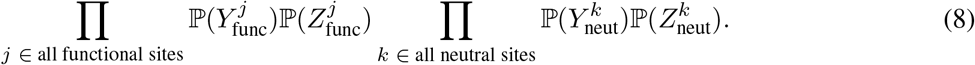

It is worth noting that the MK regression currently can only be applied to a nucleotide site where all the potential mutations have similar effects, such as 0D and 4D sites in coding regions.

I estimate model parameters (*β*_0_, *β*_*i*_, *γ*_0_, *γ*_*i*_, *P*_neut_, and *D*_neut_) by maximizing the logarithm of the MK-based likelihood function (equation 8). Also, I estimate the standard errors of parameters using the observed Fisher information matrix and compute the *p*-values of estimated parameters using the (two-tailed) Wald test.

### Simulation experiments

In each simulation run, I first sampled genomic features 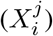 at 10 Mb functional sites from a bivariate normal distribution. I set the means and variances of the bivariate normal distribution to 0 and 1, respectively, and varied its correlation coefficient from 0 to 0.6 to examine the performance of the MK regression with respect to various degrees of correlation between features. Given genomic features, I generated divergence and polymorphism data following the likelihood function of the MK regression. Specifically, I generated 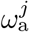, the rate of adaptive evolution at each functional site *j*, using equation 1 in the MK regression model. Also, I generated 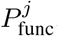, the polymorphic rate at each functional site *j*, using equation 2 in the MK regression. Finally, I generated binary response variables of polymorphism and divergence at 10 Mb neutral sites (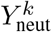 and 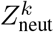) and 10 Mb functional sites (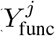 and 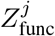) using equations 3, 4, 5, and 7 in the MK regression.

I performed two simulation experiments each of which consisted of four sets of synthetic data. In the first experiment, I set *β*_0_ = −2, *β*_1_ = 1, *β*_2_ = 0, *γ*_0_ = −8, and sampled *γ*_1_ and *γ*_2_ from a normal distribution with a mean of 0 and a standard deviation of 0.5 in each simulation run. These parameters were chosen to ensure that genome-wide levels of polymorphism and divergence are comparable between synthetic and empirical data. I generated four sets of synthetic data with different correlation coefficients (0, 0.2, 0.4, and 0.6) between features in the aforementioned bivariate normal distribution. In each dataset, I performed 10 independent simulation runs using the method described in the previous paragraph. In the second experiment, I generated synthetic data using the same procedure but replaced *β*_2_ with -0.2.

### Polymorphism and divergence data

Throughout this work, I focused on analyzing 0D and 4D sites within previously defined callable regions on autosomal chromosomes in the panTro4 reference genome (de Manuel *et al*., 2016). I obtained whole genome sequencing (WGS) based genotypes of 18 central chimpanzee (*Pan troglodytes troglodytes*) individuals from de Manuel *et al*. (2016). I filtered out all multiallelic sites and sites with missing genotypes. Then, I filtered out SNPs without a high-confidence ancestral allele (see below) and SNPs with a derived allele frequency below 0.5.

I obtained high-confidence chimpanzee ancestral alleles from Ensembl release 75 (Herrero *et al*., 2016; Yates *et al*., 2020). Ensembl reconstructed chimpanzee ancestral alleles using a phylogenetic approach, and defined that an ancestral allele was of high confidence if it was identical to both the human reference allele and the reconstructed allele in the human-chimpanzee-macaque ancestor. To annotate interspecies divergence in the chimpanzee lineage, I compared the chimpanzee ancestral allele with the reference allele in panTro4 at each monoallelic site.

### Estimation of *α* and *ω*_a_

Based on the aforementioned polymorphism and divergence data in chimpanzee autosomal genes, I computed *d*_*n*_, *p*_*n*_, *d*_*s*_, and *p*_*s*_, which are the proportions of 0D sites with divergence, 0D sites with polymorphism, 4D sites with divergence, and 4D sites with polymorphism, respectively. Each of these proportions was calculated as the ratio of the number of sites with divergence/polymorphism to the total number of sites. I estimated the proportion of adaptive substitutions as (Charlesworth, 1994; Smith and Eyre-Walker, 2002)

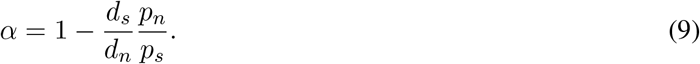

Similarly, given the polymorphism and divergence data in a gene group of interest, I estimated the relative rate of adaptive substitutions with respect to neutral evolution as (Smith and Eyre-Walker, 2002; Booker *et al*., 2017; Fraïsse *et al*., 2019)

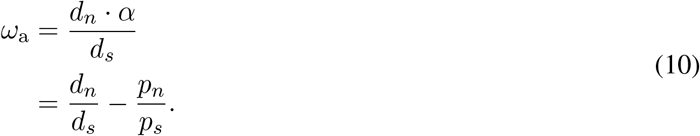

I also used equation 10 to estimate *ω*_a_ for synthetic data, where *d*_*n*_, *p*_*n*_, *d*_*s*_, and *p*_*s*_ were interpreted as the proportions of simulated functional sites with divergence, simulated functional sites with polymorphism, simulated neutral sites with divergence, and simulated neutral sites with polymorphism, respectively.

### Genomic features

Because gene annotations and functional genomic data were more complete and of higher quality in humans than in chimpanzees, I obtained human-based annotations of 0D sites, 4D sites, residue exposure level, gene expression level, tissue specificity, and protein-protein interactions. Then, I converted these annotations from the human reference genome to the panTro4 assembly using liftOver (Haeussler *et al*., 2019). Because of the very short divergence time between humans and chimpanzees, human-based genomic features should serve as an accurate proxy for the corresponding genomic features in chimpanzees. Specifically, I obtained the coordinates of 0D and 4D sites in the human genome from dbNSFP version 4.0 (Liu *et al*., 2013, 2016), and converted the coordinates from the original hg19 assembly to the panTro4 assembly using liftOver (Haeussler *et al*., 2019). I obtained predicted probabilities of residue exposure (PredRSAE) in the hg19 assembly from SNVBox (Wong *et al*., 2011). I obtained human-based consensus RNA expression levels across 62 tissues from the Human Protein Atlas version 19.3 (Uhlen *et al*., 2015). For each human protein-coding gene, I computed its (mean) expression level,

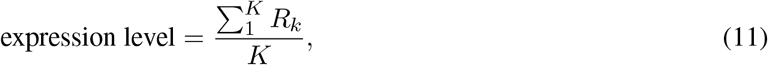

in which *R*_*k*_ is the gene’s consensus RNA expression level in tissue *k* and *K* is the total number of tissues. I computed the tissue specificity (tau) of each human gene,

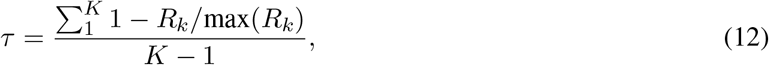

in which max(*R*_*k*_) is the gene’s maximum expression level across all tissues (Yanai *et al*., 2005). Also, I computed an alternative metric of tissue specificity (the negative value of Hg; Kryuchkova-Mostacci and Robinson-Rechavi 2017),

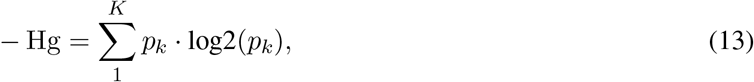

Where 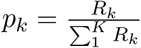 is the normalized expression level of a gene in tissue *k*. I obtained 2,274 genes with biased or enriched expression in digestive organs, including intestine, liver, pancreas, salivary gland, and stomach, from the Human Protein Atlas (Uhlen *et al*., 2015). I obtained human protein-protein interaction data from HuRI (Luck *et al*., 2020) and computed each gene’s PPI degree, *i*.*e*., the total number of unique interaction partners. Also, I obtained 2,220 human metabolic genes involved in one or more curated metabolic pathways and 3,400 human genes involved in one or more immune system pathways from MSigDB release 7.1 (Subramanian *et al*., 2005; Liberzon *et al*., 2011). Finally, I converted human-based annotations of residue exposure level, gene expression level, tissue specificity, PPI degree, metabolic genes, and immune system genes from the hg19 assembly to the panTro4 assembly using liftOver (Haeussler *et al*., 2019).

I used chimpanzee-based data to build a map of local recombination rates and a map of local mutation rates. Specifically, I obtained a fine-scale chimpanzee genetic map from panMap (Auton *et al*., 2012) and converted the data from panTro2 to panTro4 using liftOver. Then, I constructed a map of local recombination rates by averaging the recombination rates from the chimpanzee genetic map with a 1Mb non-overlapping sliding window. Also, I utilized interspecies divergence in the chimpanzee lineage to construct a map of local mutation rates in the panTro4 assembly. To do so, I converted putatively neutral regions defined in Huang (2020) from hg19 to panTro4 using liftOver. Then, I computed the density of chimpanzee-specific substitutions in putatively neutral regions using a 100 Kb non-overlapping sliding window, which was used as a proxy of local mutation rates.

### Estimating the effects of genomic features on the rate of adaptation

I fit the MK regression to one genomic feature at a time, which I named as the simple MK regression. To evaluate if a logarithmic transformation can improve model fitting, I carried out two analyses for each feature. In the first analysis, I standardized the feature by subtracting its mean and dividing by its standard deviation, and then fit the simple MK regression to the standardized feature. In the second analysis, I calculated the logarithm of each feature, standardized the output, and then fit the simple MK regression to the transformed data. Because the logarithm of PPI degree is undefined if the PPI degree is equal to 0, I added a pseudo-count of 1 to the PPI degree before the logarithmic transformation. I computed the log likelihood of the simple MK regression in each analysis, and used the transformation with a higher log likelihood for each feature throughout this work. I also fit the MK regression to all the features simultaneously, which I named as the multiple MK regression.

### Statistical matching analysis

I used statistical matching algorithms to estimate the effects of PPI degree, gene expression level, and tissue specificity after adjusting for other correlated genomic features. To estimate the effect of PPI degree, I stratified protein-coding genes into two groups based on PPI degree. The first group consisted of 1,556 genes with 10 or more interaction partners (PPI degree ≥ 10) while the second group consisted of 8,471 genes with 1 or less interaction partners (PPI degree ≤ 1). Here I chose a 10-fold difference in PPI degree between the two gene groups, because a larger difference in PPI degree led to significantly smaller gene groups, whereas a smaller difference in PPI degree may attenuate the potential difference in *ω*_a_ between the two gene groups. I calculated *ω*_a_ for the two groups of genes separately using equation 10, and calculated the *p*-value of the difference in *ω*_a_ between the two gene groups using a two-tailed permutation test with 1,000 resamplings. Also, I used MatchIt to match each gene from the high-PPI group with a gene of similar log expression level and tissue specificity from the low-PPI group (Ho *et al*., 2011). Then, I repeated the calculation of *ω*_a_ and *p*-value for the two groups of matched genes.

To estimate the effect of tissue specificity after controlling for gene expression level, I stratified highly expressed genes (mean expression level > 30) into two approximately equal-sized groups based on the ranking of their tissue specificity. The first gene group consisted of 368 highly expressed genes with higher tissue specificity while the second group consisted of 369 highly expressed genes with lower tissue specificity. Then, I calculated *ω*_a_ for the two groups of genes separately using equation 10, and calculated the *p*-value of the difference in *ω*_a_ using a two-tailed permutation test with 1,000 resamplings.

Finally, I estimated the effect of gene expression level after controlling for tissue specificity. I stratified tissue-specific genes (tau > 0.85) into two approximately equal-sized groups based on the ranking of their expression levels. The first group consisted of 495 tissue-specific genes with higher expression level while the second group consisted of 498 tissue-specific genes with lower expression level. I calculated *ω*_a_ for the two groups of genes separately using equation 10, and calculated the *p*-value of the difference in *ω*_a_ using a two-tailed permutation test with 1,000 resamplings.

### Reanalysis of data from Fraïsse *et al*. (2019)

I estimated the effects of gene expression level and tissue specificity on the rate of adaptive evolution in *Drosophila melanogaster* by reanalyzing data from Fraïsse *et al*. (2019). I obtained gene-level estimates of *ω*_a_, mean expression levels, tissue-by-stage specificity (tau), polymorphism data, and divergence data from http://doi.org/10.15479/at:ista:/5757. In the analysis of standard linear regression, I regressed the gene-level estimate of *ω*_a_ on the logarithm of mean expression level and the tissue-by-state specificity using the *lm* function in R (R Core Team, 2017). In the analysis of multiple MK regression, I used 0D sites and 4D sites annotated by SIFT 4G (Vaser *et al*., 2016) as functional and putatively neutral sites, respectively, and used the logarithm of mean expression level and the tissue-by-state specificity as input features. In the statistical matching analysis, I stratified 681 tissue-specific genes (tau > 0.85) into a group of 340 genes with higher expression level and a group of 341 genes with lower expression level. I calculated *ω*_a_ for the two groups of genes separately using equation 10, and calculated the *p*-value of the difference in *ω*_a_ using a two-tailed permutation test with 1,000 resamplings.

### Tissue and pathway enrichment

I downloaded annotations of tissue-enriched genes from the Human Protein Atlas version 19.3 (Uhlen *et al*., 2015). To reduce the burden of multiple testing, I focused on analyzing tissues with at least 50 tissue-enriched genes. I then analyzed the enrichment of each set of tissue-enriched genes in the 495 tissue-specific genes with high expression level, in which the 498 tissue-specific genes with low expression level were used as a background gene set. The *p*-value of each enrichment test was computed using the Fisher’s exact test and then adjusted for multiple testing with false-discovery-rate correction. Similarly, I analyzed the enrichment of Reactome pathways in the 495 tissue-specific genes with high expression level using PANTHER (Mi *et al*., 2017), in which I used the 498 tissue-specific genes with low expression level as a background gene set.

## Data availability

The MK regression model and companion data are available at https://github.com/yifei-lab/MK-regression.

## Acknowledgments

The author acknowledges David McCandlish, Xinru Zhang, Jui-Shan Lin, and Zhihan Liu for useful discussions. Research reported in this publication was supported by the National Institute of General Medical Sciences of the National Institutes of Health under Award Number R35GM142560 and by startup funds from Pennsylvania State University. The content is solely the responsibility of the author and does not necessarily represent the official views of the National Institutes of Health.

