## Supplementary Figures and Tables. for "Dissecting genomic determinants of positive selection with an evolution-guided regression model"

Yi-Fei Huang[1,2]

[1]Department of Biology, Pennsylvania State University,
University Park, PA 16802, USA

[2]Huck Institutes of the Life Sciences, Pennsylvania State University,
University Park, PA 16802, USA

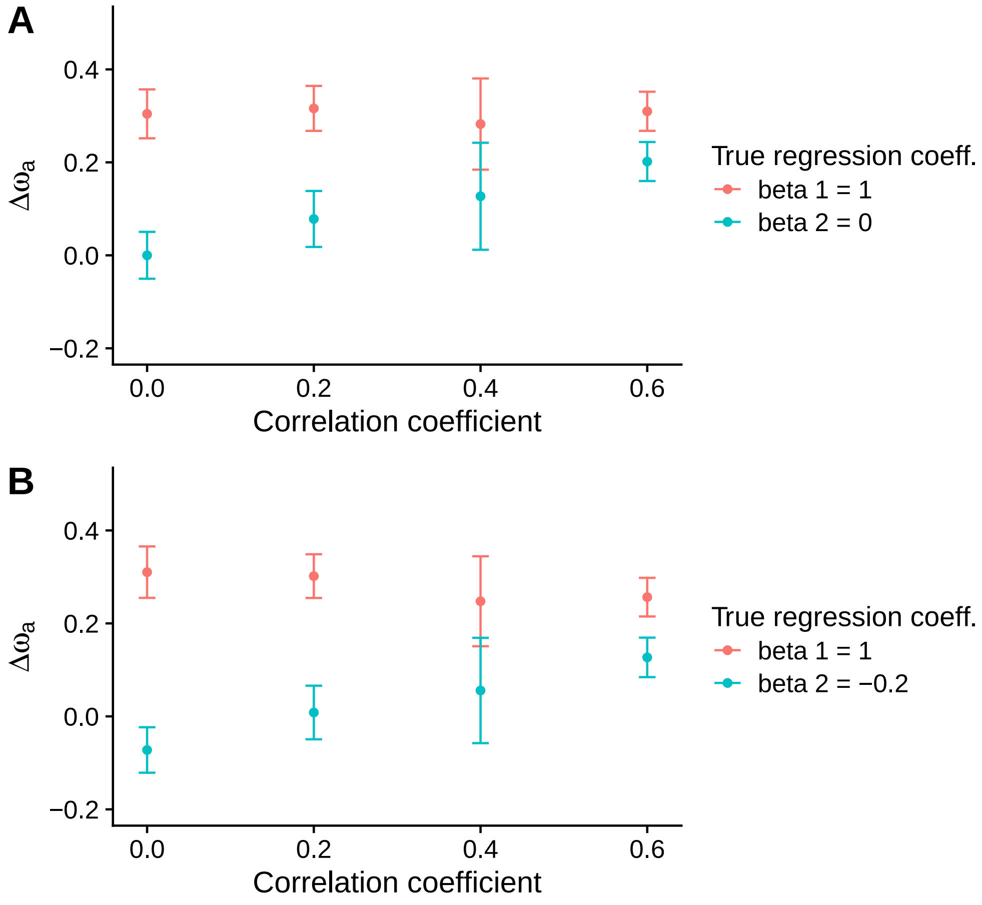


Supplementary Fig. 1: Performance of a previous MK-based method (Smith and Eyre-Walker, 2002; Fraïsse *et al.*, 2019) in synthetic data. $\Delta\omega_a$ indicates the difference in estimated $\omega_a$ between the top half of functional sites with higher feature values and the bottom half with lower feature values. In each plot, dots and error bars indicate the means and the two-fold standard deviations of $\Delta\omega_a$ across 10 independent replicates, respectively. (A) The true coefficients are $\beta_1 = 1$ and $\beta_2 = 0$. (B) The true coefficients are $\beta_1 = 1$ and $\beta_2 = -0.2$.

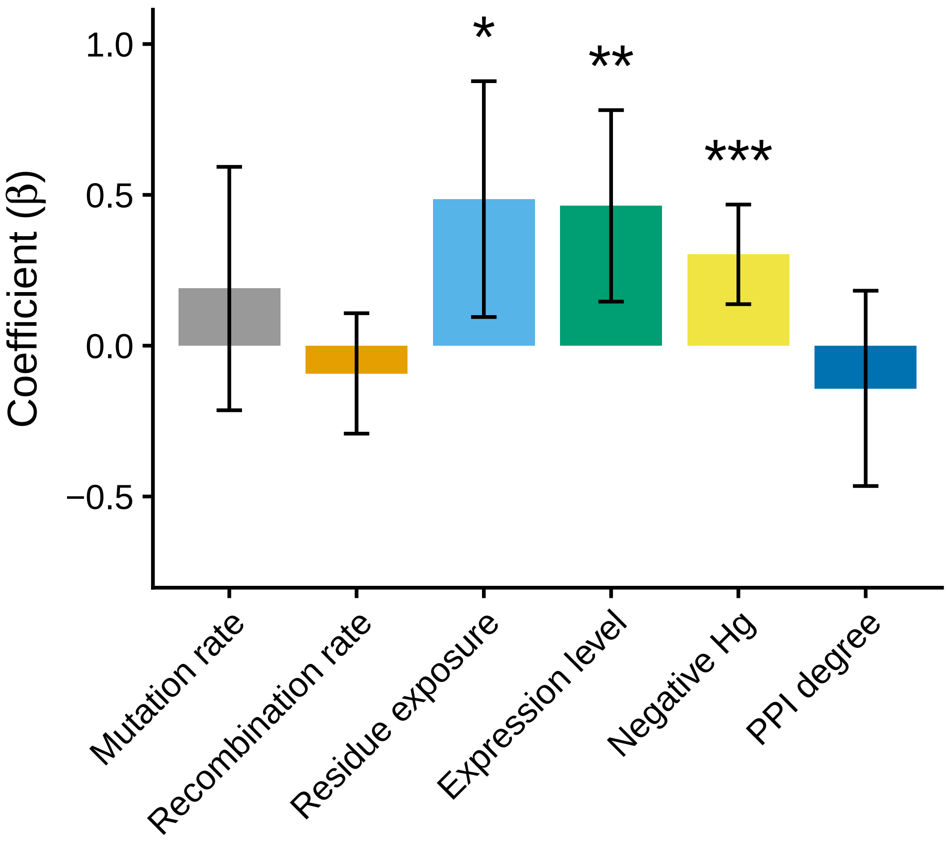


Supplementary Fig. 2: Multiple MK regression analysis. This analysis is similar to that in Fig. 5 but uses the negative value of Hg as an alternative metric of tissue specificity. Error bars indicate 95% confidence intervals while one, two, and three asterisks indicate $0.01 \leq p\text{-value} < 0.05$, $0.001 \leq p\text{-value} < 0.01$, and $p\text{-value} < 0.001$, respectively.

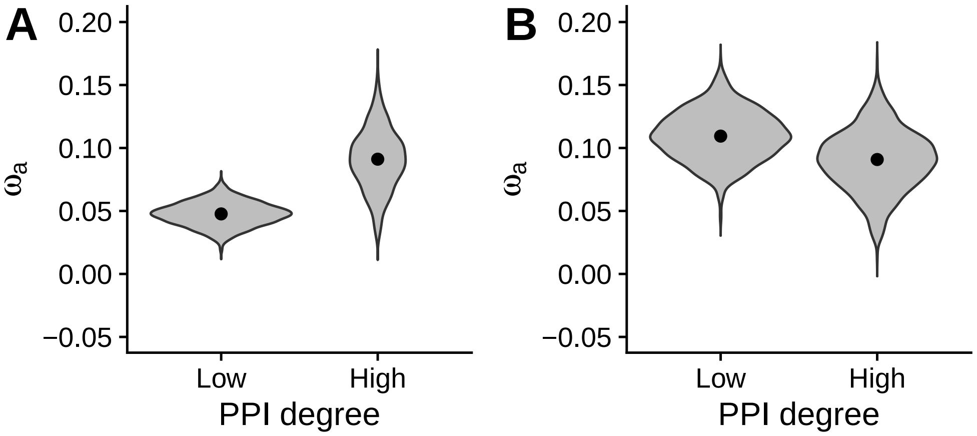


Supplementary Fig. 3: Propensity score matching analysis. (A) Estimates of $\omega_a$ in 1,556 high PPI-degree genes and 8,471 low PPI-degree genes. (B) Estimates of $\omega_a$ in 1,556 high PPI-degree genes and 1,556 low PPI-degree genes with matched log gene expression level and tissue specificity.

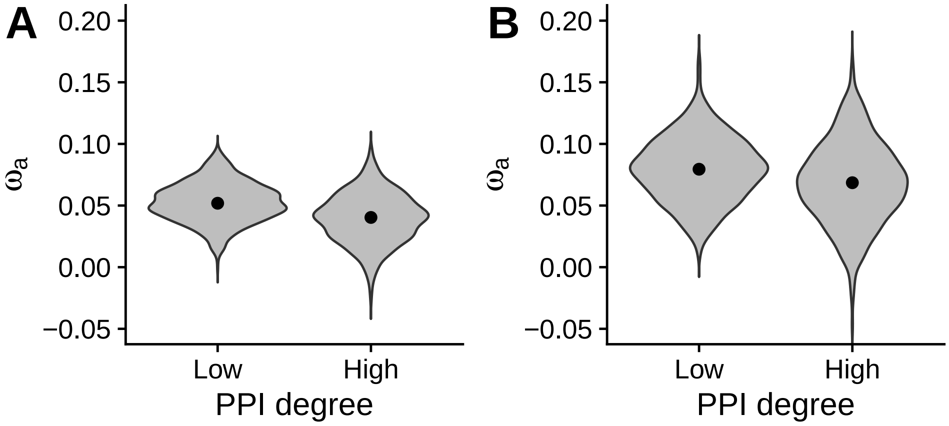


Supplementary Fig. 4: Propensity score matching analysis with alternative cutoffs for PPI degree. (A) Estimates of $\omega_a$ in 2,469 high PPI-degree genes (PPI degree $\geq$ 5) and 2,469 low PPI-degree genes (PPI degree $\leq$ 1) with matched log gene expression level and tissue specificity. (B) Estimates of $\omega_a$ in 893 high PPI-degree genes (PPI degree $\geq$ 20) and 893 low PPI-degree genes (PPI degree $\leq$ 1) with matched log gene expression level and tissue specificity.

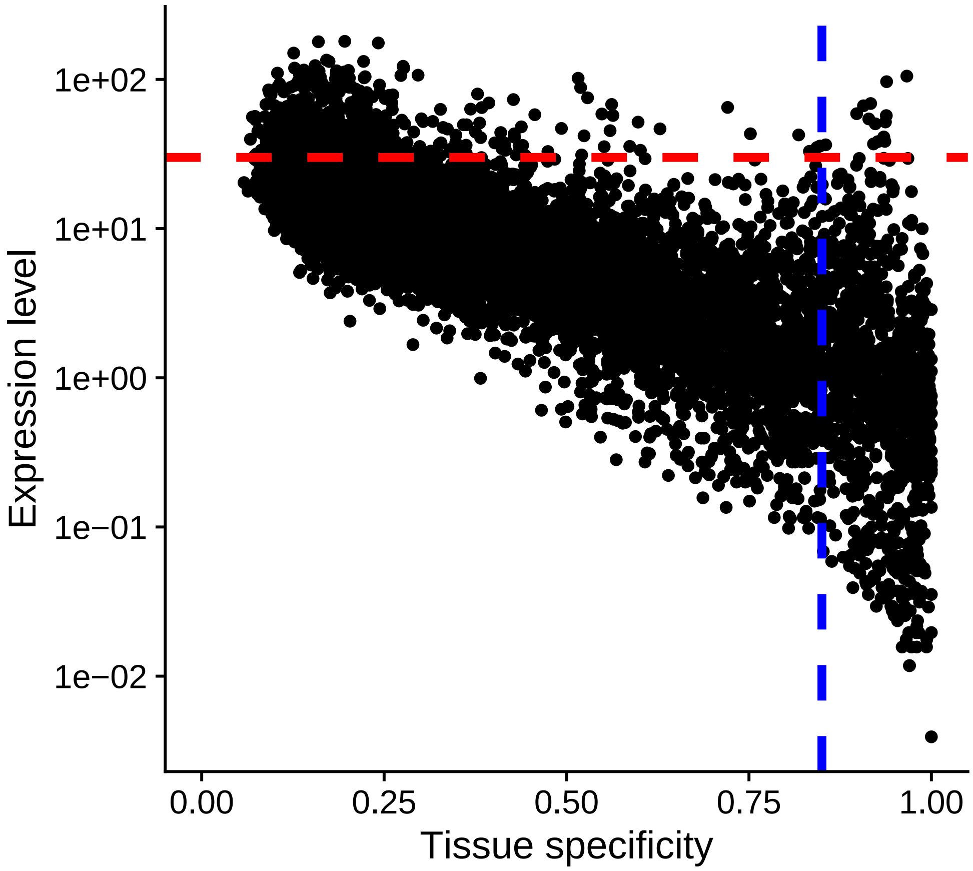


Supplementary Fig. 5: Correlation between gene expression level and tissue specificity. The red dashed line indicates the threshold for highly expressed genes (expression level $> 30$). The blue dashed line indicates the threshold for tissue-specific genes (tau $> 0.85$).

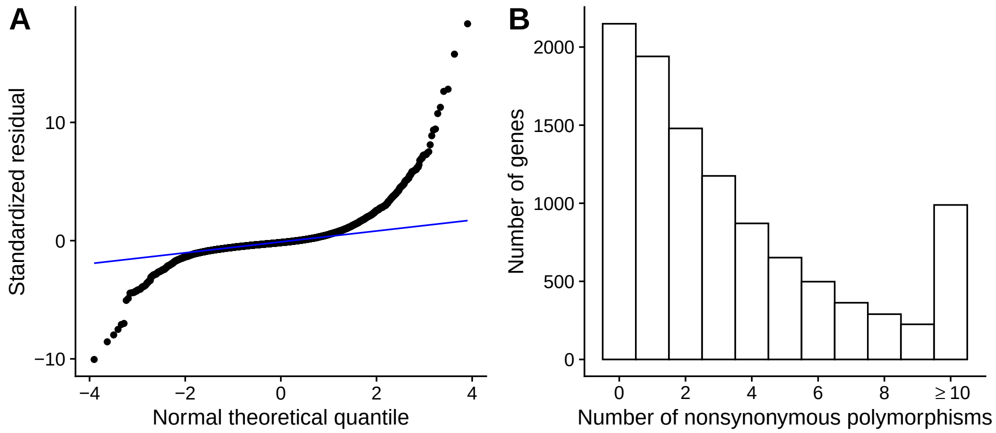


Supplementary Fig. 6: Model misspecification of the standard linear regression in the inference of adaptive evolution in *Drosophila melanogaster*. (A) The quantile-quantile plot of standardized residuals from the standard linear regression versus theoretical quantiles from a normal distribution. In the standard linear regression, gene-level estimate of $\omega_{\mathrm{a}}$ is used as a response variable, whereas log gene expression level and tissue specificity are used as covariates. The departure of data points from the blue line indicates that the residuals do not follow a normal distribution. (B) The distribution of the number of nonsynonymous polymorphisms across protein-coding genes in *Drosophila melanogaster*. The number of nonsynonymous polymorphisms is truncated at 10.

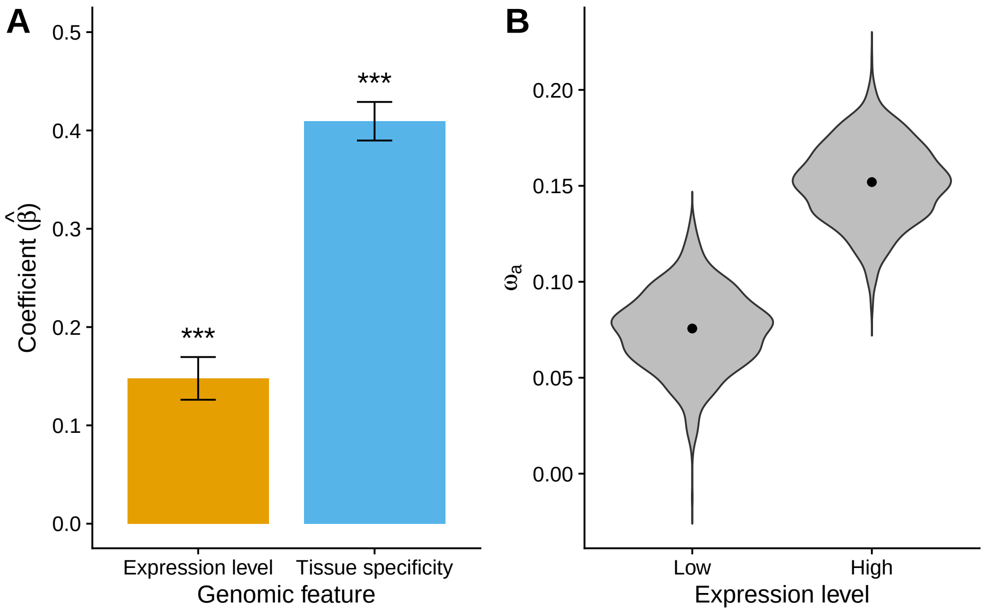


Supplementary Fig. 7: Effects of gene expression level and tissue specificity on the rate of adaptive evolution in *Drosophila melanogaster*. (A) Estimated coefficients in the Multiple MK regression. Error bars indicate 95% confidence intervals while three asterisks indicate $p$-value $< 0.001$. (B) Estimates of $\omega_a$ in 341 tissue-specific genes with low expression level and 340 tissue-specific genes with high expression level. In each violin plot, dots indicate point estimates of $\omega_a$ while violins depict the distributions of $\omega_a$ from a gene-based bootstrapping analysis with 1,000 resamplings.

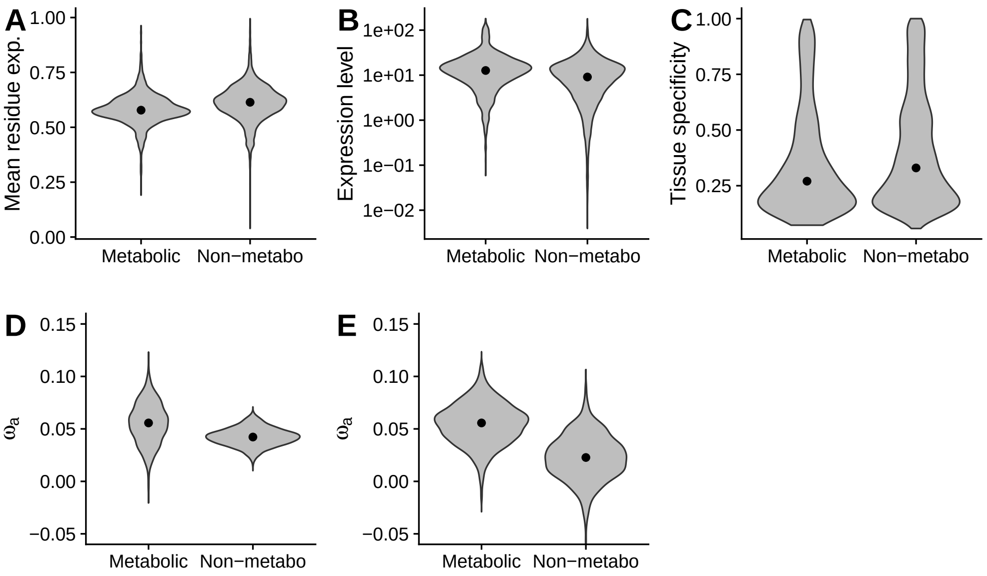


Supplementary Fig. 8: Propensity score matching analysis of metabolic genes. (A) Distributions of mean residue exposure level in metabolic and non-metabolic genes. (B) Distributions of gene expression level in metabolic and non-metabolic genes. (C) Distributions of tissue specificity (tau) in metabolic and non-metabolic genes. (D) Estimates of $\omega_a$ without controlling for residue exposure level, gene expression level, and tissue specificity. (E) Estimates of $\omega_a$ after controlling for residue exposure level, gene expression level, and tissue specificity.

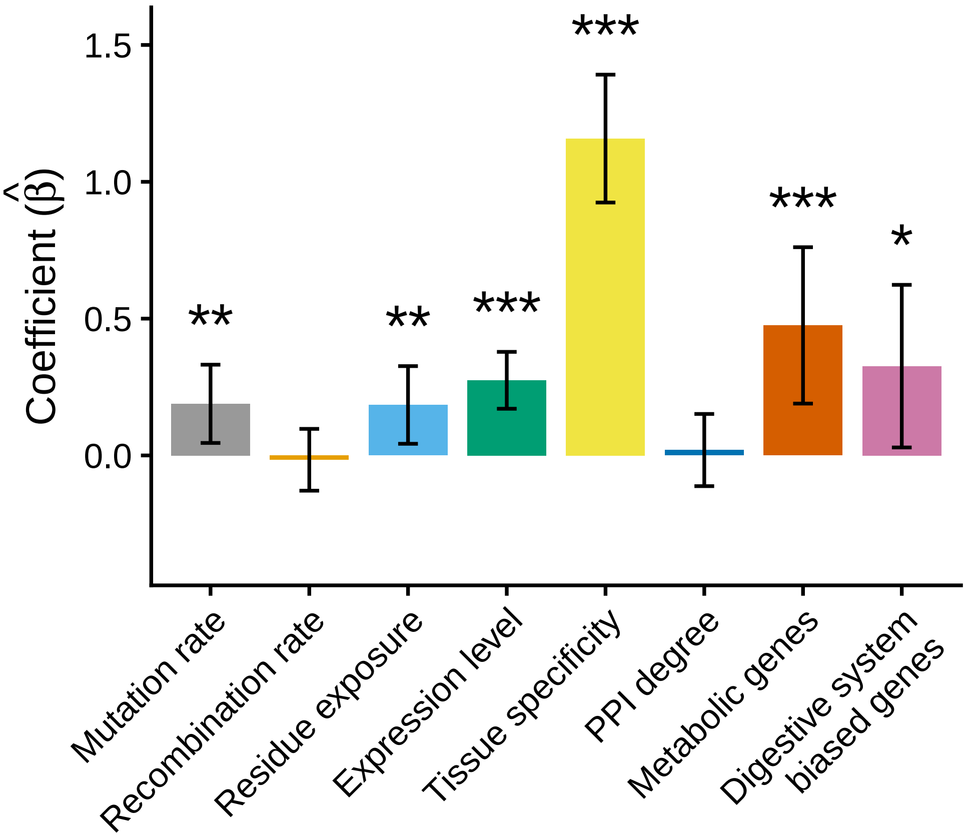


Supplementary Fig. 9: Multiple MK regression analysis of digestive system-biased genes. This analysis includes all features in Fig. 5 and a new binary feature indicating whether each 0D site is located in one of the 2,274 genes with biased or enriched expression in digestive organs, including intestine, liver, pancreas, salivary gland, and stomach, from the Human Protein Atlas (Uhlen *et al.*, 2015). Error bars indicate 95% confidence intervals while one, two, and three asterisks indicate $0.01 \leq p\text{-value} < 0.05$, $0.001 \leq p\text{-value} < 0.01$, and $p\text{-value} < 0.001$, respectively.

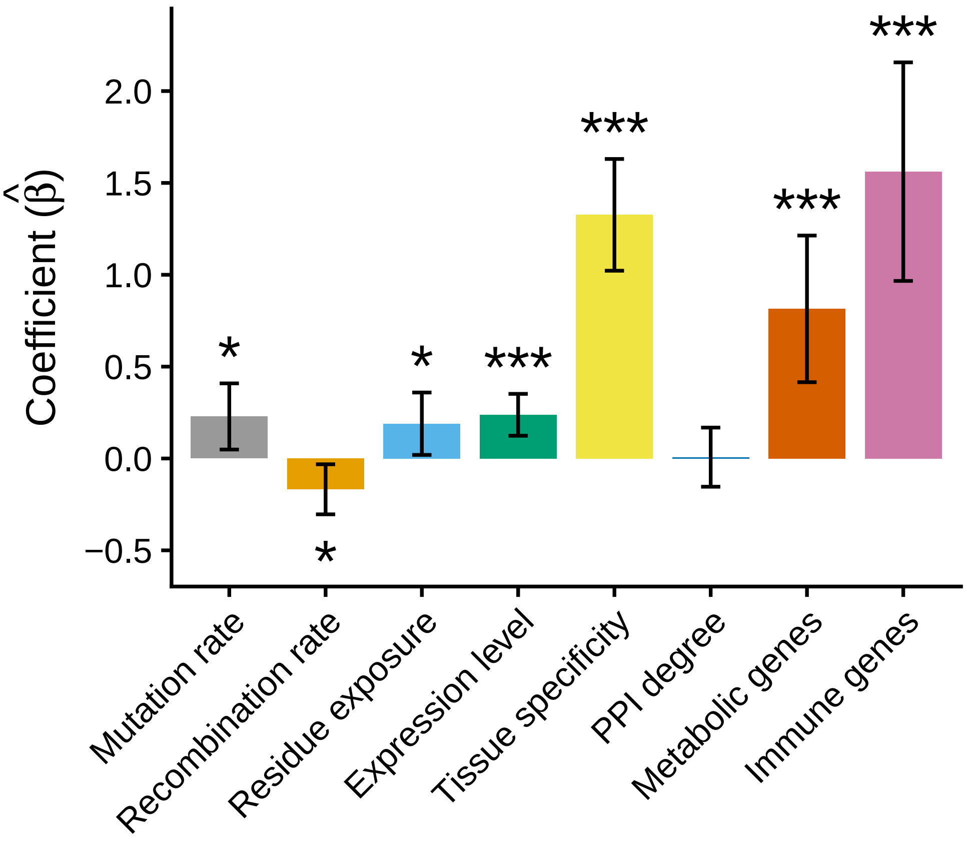


Supplementary Fig. 10: Multiple MK regression analysis of immune system genes. This analysis includes all features in Fig. 5 and a new binary feature indicating whether each 0D site is located in one of the 3,400 immune system genes from MSigDB (Subramanian *et al.*, 2005; Liberzon *et al.*, 2011). Error bars indicate 95% confidence intervals while one, two, and three asterisks indicate $0.01 \leq p$-value $< 0.05$, $0.001 \leq p$-value $< 0.01$, and $p$-value $< 0.001$, respectively.

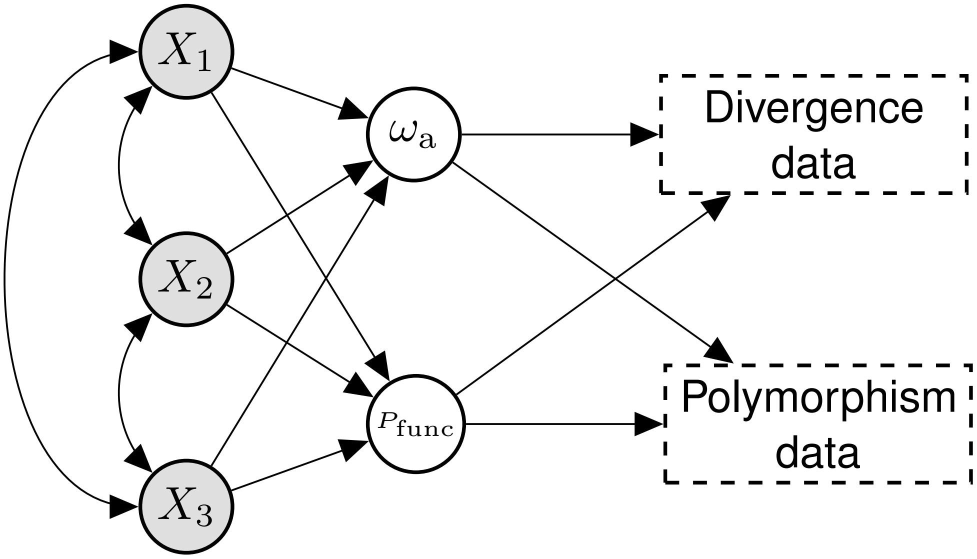


Supplementary Fig. 11: An example of the causal graph assumed in the multiple MK regression. $X_1$, $X_2$, and $X_3$ represent three genomic features at a functional site. $\omega_a$ and $P_{func}$ indicate adaptation rate and polymorphic rate at a functional site, respectively. Single-headed arrows indicate causal relationships between variables, whereas double-headed arrows indicate correlations between genomic features. The correlation between two genomic features might arise through an unspecified causal effect of one feature on the other feature, other genomic features that causally influence the two features (common causes), or both.

Supplementary Table 1: Fitting of the simple MK regression.

| Genomic feature | Log likelihood (original feature) | Log likelihood (logarithmic feature) |
|---|---|---|
| Mutation rate | −148 452.785 | −148 421.085 |
| Recombination rate | −148 568.492 | −148 568.127 |
| Residue exposure | −148 373.595 | −148 317.301 |
| Expression level | −147 768.446 | −147 704.267 |
| Tissue specificity | −147 757.508 | −147 796.508 |
| PPI degree | −148 565.259 | −148 552.632 |

Supplementary Table 2: Estimates of regression coefficients in the simple MK regression. $\hat{\beta}$ indicates an estimated regression coefficient for adaptation rate ($\omega_a$) whereas $\hat{\gamma}$ indicates an estimated regression coefficient for polymorphic rate ($P_{func}$).

| Genomic feature | Coefficient ($\hat{\beta}$) | $p$-value ($\hat{\beta}$) | Coefficient ($\hat{\gamma}$) | $p$-value ($\hat{\gamma}$) |
|---|---|---|---|---|
| Mutation rate | 0.383 | $3.372 \cdot 10^{-12}$ | 0.124 | $6.641 \cdot 10^{-15}$ |
| Recombination rate | −0.084 | 0.447 | 0.025 | 0.301 |
| Residue exposure | 0.484 | 0.002 | 0.232 | $2.024 \cdot 10^{-27}$ |
| Expression level | −0.310 | $3.534 \cdot 10^{-10}$ | −0.342 | $1.034 \cdot 10^{-236}$ |
| Tissue specificity | 0.157 | 0.622 | 0.382 | $1.577 \cdot 10^{-24}$ |
| PPI degree | 0.460 | $3.178 \cdot 10^{-4}$ | −0.140 | $3.501 \cdot 10^{-09}$ |

Supplementary Table 3: Estimates of regression coefficients in the multiple MK regression. $\hat{\beta}$ indicates an estimated regression coefficient for adaptation rate ($\omega_a$) whereas $\hat{\gamma}$ indicates an estimated regression coefficient for polymorphic rate ($P_{func}$).

| Genomic feature | Coefficient ($\hat{\beta}$) | $p$-value ($\hat{\beta}$) | Coefficient ($\hat{\gamma}$) | $p$-value ($\hat{\gamma}$) |
|---|---|---|---|---|
| Mutation rate | 0.155 | 0.018 | 0.121 | $3.239 \cdot 10^{-17}$ |
| Recombination rate | −0.038 | 0.493 | 0.015 | 0.298 |
| Residue exposure | 0.188 | $7.031 \cdot 10^{-3}$ | 0.307 | $1.939 \cdot 10^{-57}$ |
| Expression level | 0.347 | $6.674 \cdot 10^{-12}$ | −0.405 | $2.079 \cdot 10^{-58}$ |
| Tissue specificity | 1.168 | $6.394 \cdot 10^{-31}$ | −0.117 | $4.443 \cdot 10^{-4}$ |
| PPI degree | 0.002 | 0.975 | −0.013 | 0.389 |

Supplementary Table 4: Kendall's correlation coefficients between genomic features.

| | Residue exposure | Recomb. rate | Mutation rate | Expression level | Tissue specificity | PPI degree |
|---|---|---|---|---|---|---|
| Residue exposure | 1.000 | −0.003 | −0.006 | 0.018 | −0.025 | 0.020 |
| Recomb. rate | −0.003 | 1.000 | 0.022 | 0.000 | 0.006 | 0.013 |
| Mutation rate | −0.006 | 0.022 | 1.000 | −0.053 | 0.074 | −0.015 |
| Expression level | 0.018 | 0.000 | −0.053 | 1.000 | −0.616 | 0.078 |
| Tissue specificity | −0.025 | 0.006 | 0.074 | −0.616 | 1.000 | −0.071 |
| PPI degree | 0.020 | 0.013 | −0.015 | 0.078 | −0.071 | 1.000 |

Supplementary Table 5: Estimation of the effect of tissue specificity (tau) after controlling for gene expression level. Genes are divided into deciles based on their expression levels. In each decile, genes are further divided into two equal-sized subgroups by the ranking of their tissue specificity. A two-tailed permutation test is employed to calculate *p*-values.

| Decile | $\omega_a$ in the subgroup with higher tissue specificity | $\omega_a$ in the subgroup with lower tissue specificity | *p*-value |
|---|---|---|---|
| 1 | 0.0712 | 0.0477 | 0.865 |
| 2 | 0.0599 | 0.167 | 0.147 |
| 3 | 0.0501 | 0.0555 | 0.944 |
| 4 | 0.0170 | -0.0259 | 0.467 |
| 5 | 0.0149 | 0.0580 | 0.430 |
| 6 | 0.0141 | 0.0642 | 0.303 |
| 7 | -0.00424 | 0.0587 | 0.154 |
| 8 | -0.0183 | 0.0422 | 0.146 |
| 9 | 0.0640 | 0.0439 | 0.555 |
| 10 | 0.0975 | -0.00499 | 0.023 |

Supplementary Table 6: Estimation of the effect of gene expression level after controlling for tissue specificity (tau). Genes are divided into deciles based on their tissue specificity. In each decile, genes are further divided into two equal-sized subgroups by the ranking of their expression level. A two-tailed permutation test is employed to calculate *p*-values.

| Decile | $\omega_a$ in the subgroup with higher expression level | $\omega_a$ in the subgroup with lower expression level | *p*-value |
|---|---|---|---|
| 1 | 0.00515 | 0.0172 | 0.754 |
| 2 | 0.0399 | 0.102 | 0.131 |
| 3 | 0.0282 | 0.0683 | 0.393 |
| 4 | 0.0175 | 0.0703 | 0.233 |
| 5 | -0.00652 | 0.00356 | 0.816 |
| 6 | -0.0220 | -0.0566 | 0.582 |
| 7 | 0.0432 | 0.151 | 0.061 |
| 8 | -0.0664 | 0.127 | 0.017 |
| 9 | 0.0366 | 0.00712 | 0.761 |
| 10 | 0.303 | 0.013264 | 0.010 |

Supplementary Table 7: Estimates of regression coefficients in the multiple MK regression. This analysis includes a new binary feature indicating whether each 0D site is located in a metabolic gene. $\hat{\beta}$ indicates an estimated regression coefficient for adaptation rate ($\omega_a$) whereas $\hat{\gamma}$ indicates an estimated regression coefficient for polymorphic rate ($P_{func}$).

| Genomic feature | Coefficient ($\hat{\beta}$) | $p$-value ($\hat{\beta}$) | Coefficient ($\hat{\gamma}$) | $p$-value ($\hat{\gamma}$) |
|---|---|---|---|---|
| Mutation rate | 0.172 | 0.011 | 0.120 | $1.878 \cdot 10^{-18}$ |
| Recombination rate | $-0.022$ | 0.693 | 0.012 | 0.362 |
| Residue exposure | 0.184 | $7.395 \cdot 10^{-3}$ | 0.303 | $1.703 \cdot 10^{-62}$ |
| Expression level | 0.314 | $1.103 \cdot 10^{-10}$ | $-0.379$ | $2.685 \cdot 10^{-51}$ |
| Tissue specificity | 1.197 | $7.882 \cdot 10^{-28}$ | $-0.097$ | 0.004 |
| PPI degree | 0.010 | 0.872 | $-0.016$ | 0.259 |
| Metabolic gene | 0.537 | $1.075 \cdot 10^{-4}$ | $-0.396$ | $1.389 \cdot 10^{-12}$ |

Supplementary Table 8: Estimates of regression coefficients in the multiple MK regression. This analysis includes a new binary feature indicating whether each 0D site is located in an immune system gene. $\hat{\beta}$ indicates an estimated regression coefficient for adaptation rate ($\omega_a$) whereas $\hat{\gamma}$ indicates an estimated regression coefficient for polymorphic rate ($P_{func}$).

| Genomic feature | Coefficient ($\hat{\beta}$) | $p$-value ($\hat{\beta}$) | Coefficient ($\hat{\gamma}$) | $p$-value ($\hat{\gamma}$) |
|---|---|---|---|---|
| Mutation rate | 0.229 | 0.011 | 0.117 | $1.025 \cdot 10^{-22}$ |
| Recombination rate | $-0.168$ | 0.014 | 0.019 | 0.107 |
| Residue exposure | 0.189 | 0.026 | 0.293 | $2.501 \cdot 10^{-79}$ |
| Expression level | 0.238 | $3.000 \cdot 10^{-05}$ | $-0.317$ | $2.610 \cdot 10^{-36}$ |
| Tissue specificity | 1.326 | $2.718 \cdot 10^{-18}$ | $-0.006$ | 0.876 |
| PPI degree | 0.007 | 0.928 | $-0.014$ | 0.226 |
| Metabolic gene | 0.814 | $4.495 \cdot 10^{-05}$ | $-0.375$ | $7.130 \cdot 10^{-15}$ |
| Immune gene | 1.561 | $1.522 \cdot 10^{-07}$ | $-0.185$ | $6.135 \cdot 10^{-05}$ |
